# Antagonistic interactions between CLAVATA receptors shape maize ear development

**DOI:** 10.1101/2025.08.04.667500

**Authors:** Penelope L. Lindsay, Fang Xu, Lei Liu, Panagiotis Boumpas, Andres Reyes, Byoung Il Je, Mari Ogawa-Ohnishi, Jarrett Man, Tara Skopelitis, Yoshikatsu Matsubayashi, Madelaine Bartlett, Shou-Ling Xu, David Jackson

## Abstract

Meristem activity is controlled by the CLAVATA (CLV) signaling pathway, which involves a suite of leucine rich receptor (LRR) receptors, receptor-like proteins and CLV- EMBRYO SURROUNDING REGION (CLE) peptide ligands. FASCIATED EAR 3 (FEA3) is a leucine rich receptor (LRR) receptor-like protein important for meristem maintenance in maize, and acts independently of canonical CLV receptors. Weak alleles of *fea3* can increase yield-related traits in maize, so understanding how FEA3 controls inflorescence development can maximize its potential as a crop improvement target. To identify FEA3’s interaction network, we used TurboID-based proximity labeling in maize meristems, and identified a putative co-receptor, BARELY ANY MERISTEM 1D (BAM1D). BAM1D and FEA3 proximity labeling proteomes shared over 100 proteins, including many signaling proteins, suggesting they feed into a common signaling pathway. *fea3* was epistatic to *bam1d* in the control of IM size, supporting the idea that FEA3 and BAM1D interact physically. However, *fea3* and *bam1d* act antagonistically, because *fea3* mutants had larger inflorescence meristems (IMs), whereas *bam1d* mutants produced smaller IMs. Together, this study demonstrates how *in vivo* TurboID-based proximity labeling clarifies complex genetic interactions between CLV receptors and expands our knowledge of downstream signaling components of CLV signaling pathways, which are largely uncharacterized. Our findings support the notion that multiple, partially overlapping CLV receptor complexes coordinately control meristem maintenance.

## Introduction

A hallmark of plant growth is extensive post-embryonic development, which allows plants to grow flexibly in response to their environment. Meristems contain stem cells and persist throughout the life of the plant to enable this flexible growth mode. Shoot and inflorescence meristems are tightly regulated by several overlapping mechanisms, including the CLAVATA (CLV) signaling pathway (Lindsay et al., 2024). In this pathway a stem cell promoting transcription factor gene, *WUSCHEL (WUS)*, is expressed in the organizing center of the meristem, and WUS protein moves into outer meristem layers to induce the expression of *CLV3/EMBRYO SURROUNDING REGION* (*CLE)* genes, which encode small, secreted peptides. CLE peptides are in turn perceived by a wide range of receptor proteins, including receptor kinases CLV1, BARELY ANY MERISTEM (BAM) 1, BAM2, and BAM3, RECEPTOR-LIKE PROTEIN KINASE 2 (RPK2), CLAVATA3-INSENSITIVE RECEPTOR KINASE (CIK) co-receptors, and receptor-like protein CLV2 (DeYoung et al., 2006; Kinoshita et al., 2010; Hu et al., 2018; Zhu et al., 2021). Perception of CLE peptides modulates the expression of *WUS*, restricting its expression to the organizing center. This feedback ensures the balance between stem cell proliferation with the production of new organs and tissues. The expansion of CLV receptor genes and their ligands has created sophisticated regulatory networks for meristem maintenance over evolutionary time (DeYoung et al., 2006; Rodriguez-Leal et al., 2019; Schlegel et al., 2021; Hirakawa, 2022; Seo et al., 2024; Zhang et al., 2024). Subtle modifications to the CLV pathway can increase yield-related traits in a broad range of plant species, so a deeper understanding of CLV receptor networks can help guide strategies for crop improvement (Fan et al., 2014; Je et al., 2016; Liu et al., 2021; Wang et al., 2021).

There are many unknowns about how receptor proteins coordinate to control stem cell proliferation. For example, the interplay between CLV1 and orthologous BAMs in the control of stem cell maintenance is complex. CLV1 and BAM1-3 appear to act antagonistically in the control of stem cell populations in Arabidopsis. *clv1* mutants have larger shoot apical meristems, but *bam* mutants have smaller meristems (DeYoung et al., 2006; DeYoung and Clark, 2008; Schlegel et al., 2021). Conversely, *clv1 bam1 2* and *clv1 bam1 2 3* mutants have a greatly enlarged meristem relative to *clv1* single mutants, suggesting that *BAMs* can compensate for a loss of *CLV1* (DeYoung and Clark, 2008; Nimchuk et al., 2015). In Arabidopsis, *BAMs* are actively derepressed to compensate for a lack of *CLV1*, but in tomato and groundcherry, *BAMs* passively compensate for loss of *CLV1* (Seo et al., 2024). *BAM1/2* and *BAM3* also act antagonistically in root meristem regulation (Zhang et al., 2024). Heterologous experiments in *N. benthamiana* have demonstrated that CLV receptors physically interact, but the biological relevance of these interactions remains obscure, potentially due to the pleiotropic role CLV receptors play in different aspects of plant development, complex genetic interactions between receptor mutants, and the lack of *in vivo* interaction data.

FASCIATED EAR 3 (FEA3) is an LRR receptor-like protein with a key role in meristem maintenance in maize (Je et al. 2016). *fea3* mutants produce fasciated ears, a result of an over-proliferating inflorescence meristem. *FEA3* is expressed below the organizing center, and in *fea3* mutants, *WUS* expression extends below the organizing center, indicating that FEA3 represses *WUS* from below, in contrast to the canonical pathway involving CLV1 and CLV2 orthologs THICK TASSEL DWARF (TD1) and FEA2 which operate in the outer layers of the meristem (Taguchi-Shiobara et al., 2001; Bommert et al., 2005). FEA3 perceives the FON2-LIKE CLE PROTEIN (FCP1) CLE peptide, but not ZmCLE7, the CLV3 ortholog. *FEA3* likely acts independently of *FEA2*, since *fea2;fea3* phenotypes are additive (Je et al., 2016).

Here we sought to define additional components of the FEA3 receptor signaling complex. We identified a maize *BAM* gene, *BAM1D*, as a potential co-receptor for FEA3, since it binds FCP1. Through analysis of fluorescent protein fusion lines, as well as *in vivo* TurboID-based proximity labeling coupled with affinity purification mass spectrometry (AP-MS), we show that FEA3 and BAM1D interact in ear primordia and shoot apices. We find that *bam1d* mutants have smaller ears, and that it interacts genetically with *fea3* in the control of ear development. We identify a significant overlap in shared interacting proteins between FEA3 and BAM1D, including proteins involved with signaling and receptor-mediated endocytosis, suggesting the receptors operate in a shared signaling pathway. Together, these results help define the signaling landscape of FEA3 and more broadly, help clarify how CLV receptors coordinate to control meristem activity.

## Results

### BAM1D binds the CLE peptide perceived by FEA3, and is co-expressed with FEA3

Since FEA3 is largely uncharacterized, we wanted to better understand its mode of action in meristem size regulation. FEA3 perceives the CLE peptide FCP1 in peptide assays (Je et al., 2016), so we asked if FEA3 binds FCP1 directly. However, Halo-tagged FEA3 did not bind radio-labeled FCP1 peptide in Halo assays (Figure S1A). Consistently, FCP1 was not predicted to interact strongly with FEA3 in AlphaFold-Multimer (iPTM=0.66), further supporting the idea that they do not interact directly (Figure S1B, C). Genetic analyses between *fea3* and *td1,* the maize ortholog of Arabidopsis CLV1, showed an additive relationship, suggesting that TD1 does not interact with FEA3 (Figure S2). However, FCP1 did bind to the Arabidopsis BARELY ANY MERISTEM 1 (AtBAM1) receptor, which led us to hypothesize that FEA3 might perceive FCP1 in a complex with a BAM-related receptor kinase (Figure S1A). We found seven *BAM* genes in maize, with four co-orthologous to *AtBAM1* and *AtBAM2*, called *ZmBAM1A*, *ZmBAM1B*, *ZmBAM1C*, and *ZmBAM1D*, and three co-orthologous to *AtBAM3*, called *ZmBAM3A*, *ZmBAM3B*, and *ZmBAM3C* (Figure S3). We examined the expression pattern of *BAM* genes in ear primordia using *in situ* hybridization. Of the seven *BAM* genes, *BAM1D* expression was most similar to *FEA3*, with strongest expression towards the base of spikelet and spikelet pair meristems (Figure 1A, Figure S4). A native promoter-driven BAM1D-YFP fusion was localized in a similar pattern, at the base of the inflorescence, spikelet and spikelet pair meristems, and in the vasculature (Figure 1B). BAM1D-YFP was detected on the plasma membrane and in the vacuole, consistent with Arabidopsis BAM proteins (Hazak et al., 2017; Schlegel et al., 2021). BAM1D-YFP was also expressed in both vegetative apices and root apical meristems, similar to FEA3 (Figure S5A, C, D). Therefore, we hypothesized that FEA3 and BAM1D may form a coreceptor-receptor pair. We indeed found that BAM1D binds FCP1 in a Halo assay (Figure 1C). Consistently, AlphaFold multimer predicted a strong interaction between BAM1D and FCP1 (iPTM=0.79), compared to a simulation between BAM1D and a scrambled CLE peptide (iPTM=0.62) (Figure 1D, E, Figure S1D, E). Indeed, FCP1 was predicted to bind BAM1D in a similar conformation and location as previous molecular docking analyses between Arabidopsis BAM receptors and CLE peptides (Roman et al., 2022).

**Figure 1.**
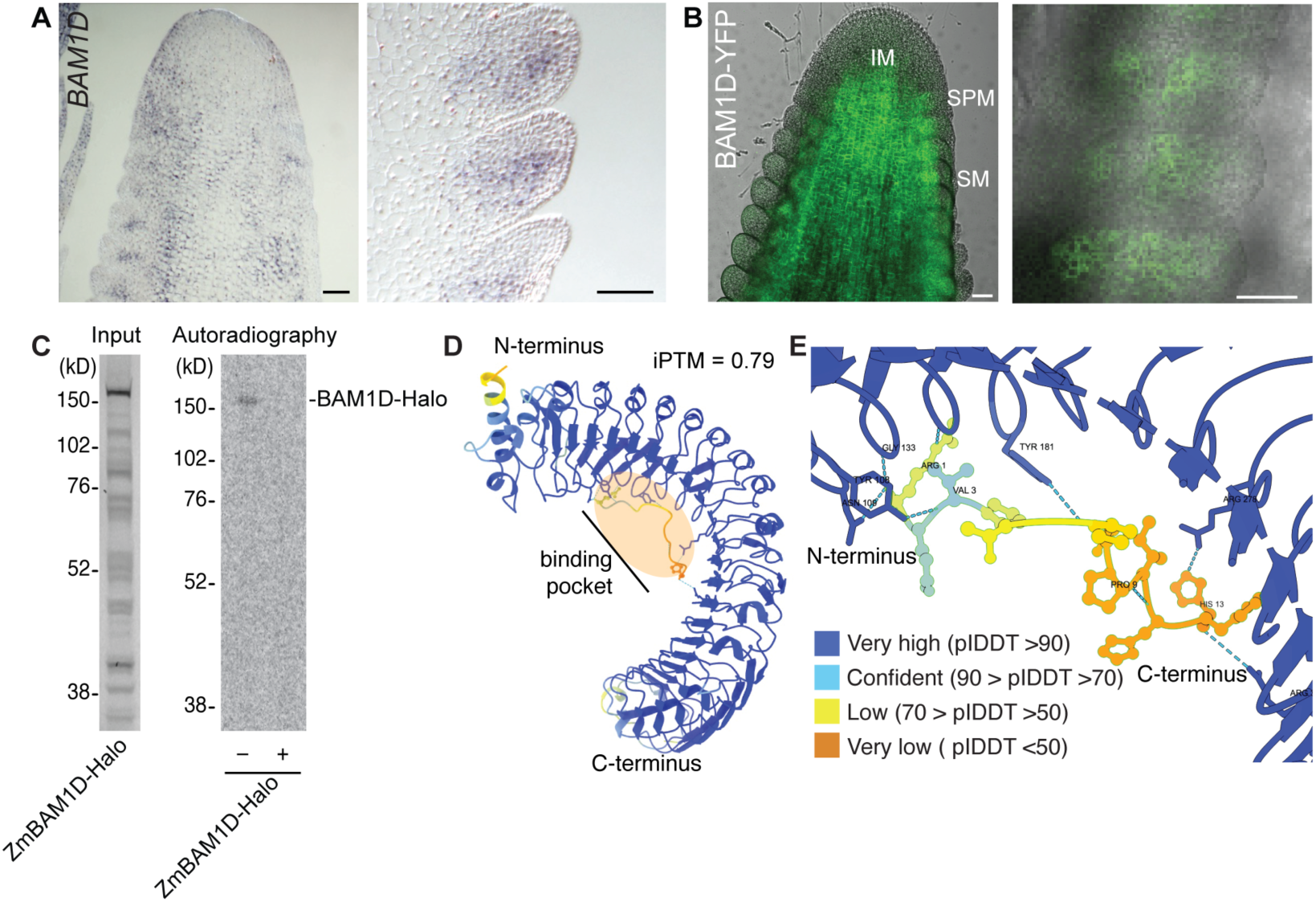
BAM1D is expressed in ear primordia and binds FCP1. (A) *in situ* hybridization of *ZmBAM1D* in ear primordia. Left panel, ear primordium overview, right panel, spikelet meristems. *ZmBAM1D* was expressed at the base of spikelet pair and spikelet meristems, and in the vasculature. (B) Localization of BAM1D-YFP in ear primordia. Left panel, ear primordium overview, right panel, spikelet meristems. BAM1D- YFP was expressed at the base of the inflorescence, spikelet pair and spikelet meristems, and vasculature. Scale bars, 50 µm. (C) FCP1 bound to ZmBAM1D-Halo in halo binding assay; binding was outcompeted by unlabeled FCP1 peptide competitor (lane marked “+”). (D) AlphaFold-Multimer prediction of BAM1D LRR and FCP1 binding interface. Predicted binding pocket highlighted in orange. BAM1D LRR domain and FCP1 interactions colored by Local Distance Difference Test (pIDDT) score, where blue indicates stronger confidence, and yellow / orange indicates weaker confidence. (E) Close-up of binding interface; dashed blue lines indicate predicted hydrogen bonds, amino acids with hydrogen bonds are labeled. Interface predicted template modeling (iPTM) score indicates the likelihood of interaction. FCP1 and BAM1D are colored by Local Distance Difference Test (pIDDT) score, where blue indicates stronger confidence, and yellow / orange indicates weaker confidence. FCP1 and BAM1D were strongly predicted to interact.

We next imaged lines expressing *BAM1D*-*YFP* and *RFP*-*FEA3,* and found overlapping expression in ear primordia, as well as in the root apical meristem (Figure 2, Figure S5C, D). Together, these data show that FEA3 and BAM1D are co-expressed in ear primordia, and suggest they perceive the same CLE peptide, FCP1.

**Figure 2.**
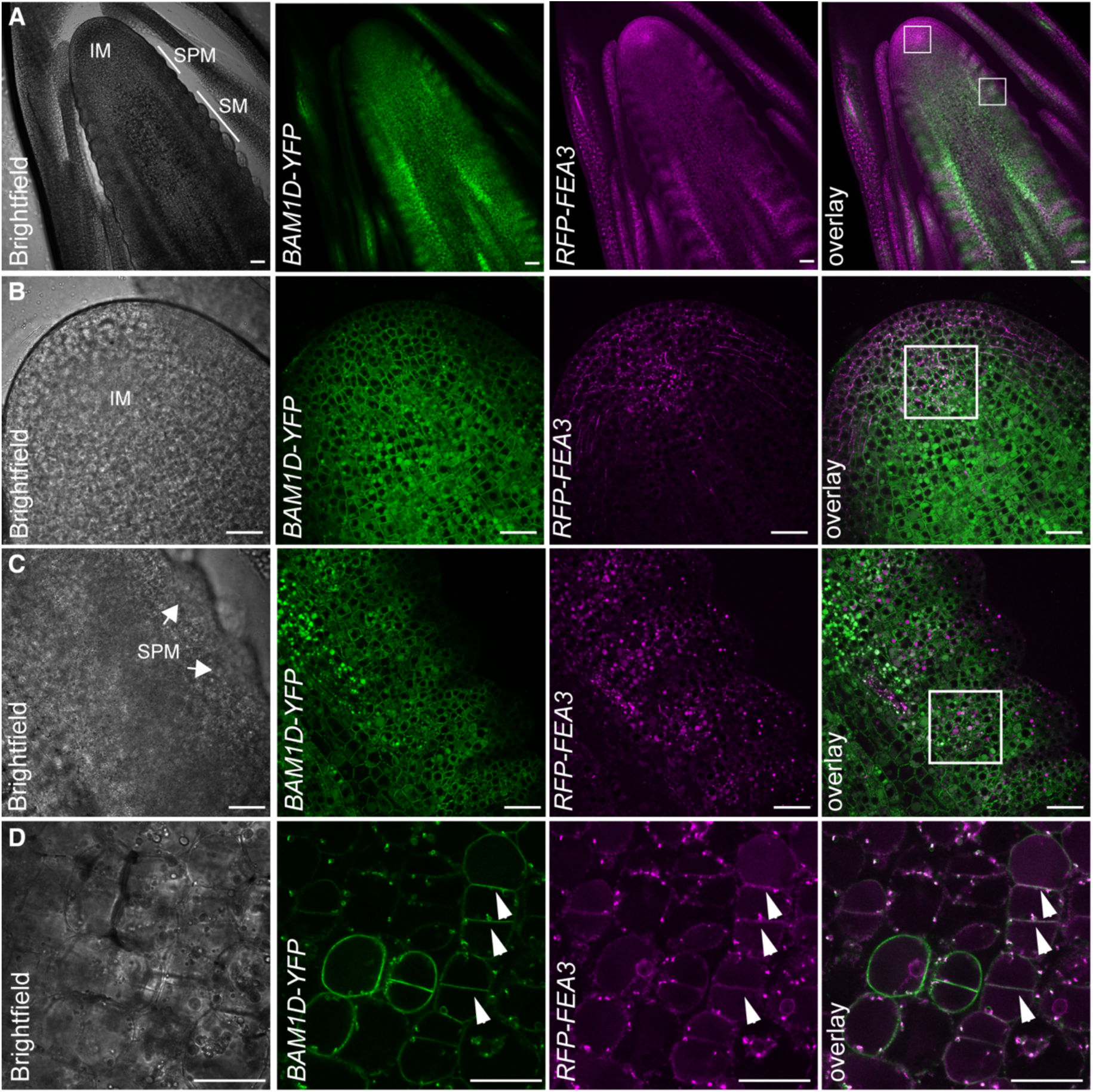
*BAM1D*-*YFP* and *RFP*-*FEA3* are co-expressed in ear primordia. (A) Overview of RFP-FEA3;BAM1D-YFP expressing ear primordia. RFP-FEA3 and BAM1D- YFP were co-expressed at the base of spikelet pair (SPM) and spikelet meristems (SM), and towards the base of the inflorescence meristem (IM). (B) Inset of IM from panel A.(C) Inset of SPM from panel A. Regions of co-expression are marked with white boxes. (D) High magnification image of the base of an RFP-FEA3;BAM1D-YFP co-expressing SPM. Co-localized signal is indicated with arrowheads. Scale bars, 50 µm.

### Proximity labeling with TurboID AP-MS reveals a FEA3-BAM1D receptor complex

To further investigate whether FEA3 and BAM1D form a receptor complex, we asked if they physically interact and if they interact with shared signaling components. To assay protein-protein interactions, we employed TurboID to label proteins in proximity to either FEA3 or BAM1D (Mair et al., 2019). We generated natively expressed FEA3-Turbo-FLAG and BAM1D-Turbo-FLAG transgenic lines (Figure 3A). We saw an increase in biotinylation over time in BAM1D-Turbo and FEA3-Turbo shoot apices, demonstrating that the TurboID fusions were able to effectively biotinylate proteins in maize (Figure S6A).

**Figure 3.**
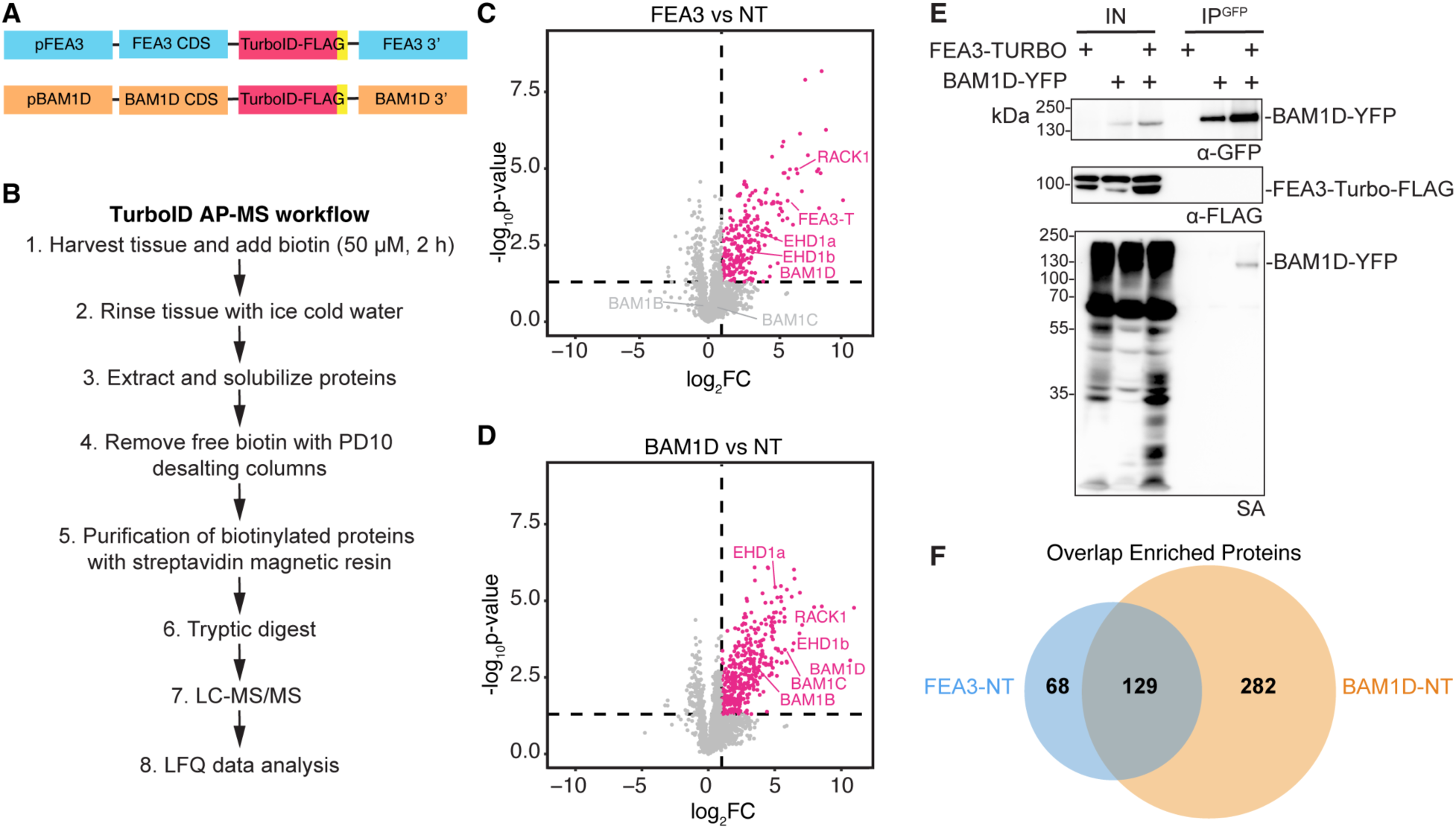
TurboID AP-MS of FEA3 and BAM1D identifies shared putative downstream signaling targets. (A) Constructs for TurboID AP-MS. p, promoter, CDS, coding sequence, 3’ downstream region. (B) TurboID AP-MS workflow. (C) Volcano plot showing FEA3-Turbo enriched proteins relative to non-transgenic control. (D) Volcano plot showing BAM1D-Turbo enriched proteins relative to non-transgenic control. (E) FEA3-Turbo biotinylated BAM1D-YFP in maize ear primordia. Bait: BAM1D-YFP. IN, input, IP, immunoprecipitated, SA, streptavidin. (F) Highly significant overlap of enriched proteins in FEA3-Turbo and BAM1D-Turbo. NT, non-transgenic.

Next, we performed TurboID affinity purification-mass spectrometry (AP-MS) on maize shoot apex tissue to identify putative downstream signaling components shared between BAM1D and FEA3, compared to a non-transgenic control (Figure 3B). Following MS analysis, we performed a principal component analysis and found that biological replicates clustered with one another (Figure S6C, D). We calculated the log2 fold change of either BAM1D-Turbo or FEA3-Turbo relative to the non-transgenic control and found significant enrichment of proteins in each condition (Figure 3C, D).

Relative to the non-transgenic control, 197 proteins were enriched in FEA3-Turbo (Figure 3C, Supplementary Table 1), including BAM1D, which was the only significantly enriched BAM receptor (Figure 3C). To validate this interaction, we crossed BAM1D-YFP plants with FEA3-Turbo plants and asked if FEA3-Turbo could biotinylate BAM1D-YFP. Indeed, FEA3-Turbo biotinylated BAM1D-YFP in ear primordia, further supporting the hypothesis that FEA3 and BAM1D are part of the same receptor complex (Figure 3E). Overrepresented GO enrichment terms in the FEA3-Turbo proximity labeling proteome included Golgi vesicle transport and endosomal transport (Figure S6E), suggesting FEA3 activity at the plasma membrane is regulated, consistent with other receptor proteins (Claus et al., 2018).

In the BAM1D-Turbo experiment, 411 proteins were enriched relative to the non-transgenic control, including additional BAM receptor kinases BAM1B and BAM1C (Figure 3D, Supplementary Table 2). Signal transduction, protein phosphorylation, endocytosis, and cell morphogenesis GO terms were enriched in BAM1D-Turbo relative to the non-transgenic control (Figure S6F). Enriched signaling proteins included auxin-related proteins TRANSMEMBRANE KINASE 1 (TMK1) and PIN-FORMED auxin transporter 1D (PIN1D), along with several MAP kinases.

Many proteins were enriched in both datasets. Specifically, 129 enriched proteins were shared between FEA3-Turbo and BAM1D-Turbo datasets, which represents a significant overlap, hypergeometric test P=2.32e^-37^ (Figure 3F, Supplementary Table 3). Shared putative interactors included proteins related to receptor-mediated endocytosis, such as C-terminal Eps15 homology domain protein (EHD1), which functions in kernel development and vegetative growth in maize and is implicated in regulating auxin dynamics (Wang et al., 2020). Signaling related proteins that were shared included the G-protein scaffold RACK1, along with 19 proteins involved in auxin signaling (Figure 3G, Supplementary Table 3) supporting the idea that FEA3 and BAM1D interact in a receptor complex in meristems.

### BAM1D perceives FCP1 and regulates maize ear size

Given our molecular data supporting FEA3 and BAM1D interaction, and that BAM1D binds FCP1, we next asked whether BAM1D operates in the same pathway as FEA3 and FCP1 to control maize ear size. To determine whether BAM1D perceives FCP1, we performed peptide treatment assays on the *bam1d-1* mutant (Figure 4A), and indeed found that the mutants were insensitive to FCP1 (Figure 4B). *bam1d-1* mutants, unlike *fea3*, were also insensitive to ZmCLE7 treatment, suggesting that BAM1D perceives a broader range of CLE peptides than FEA3 (Je et al., 2016).

**Figure 4.**
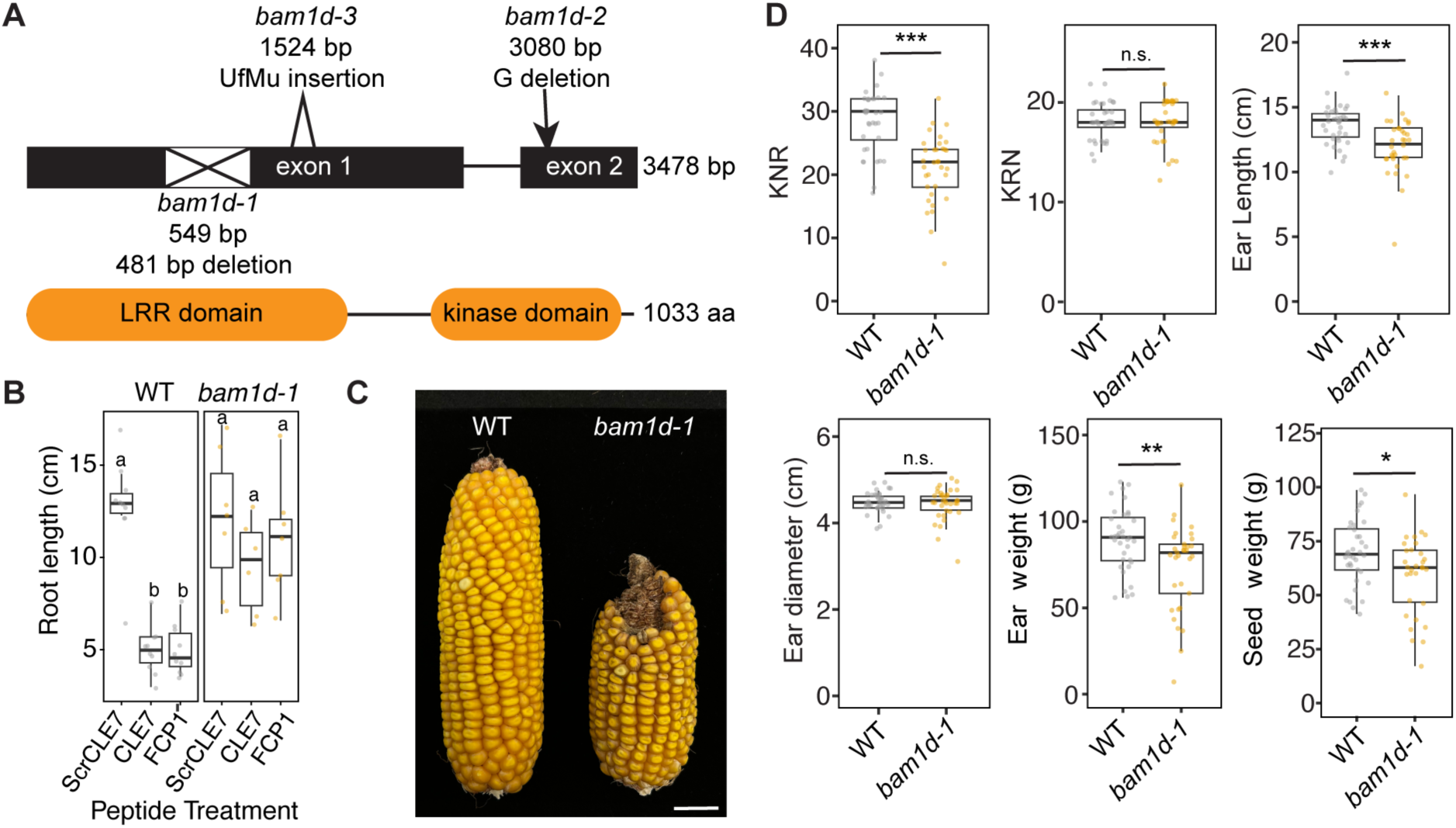
*BAM1D* regulates maize ear size. (A) Mutant alleles of *ZmBAM1D. bam1d-1* is a 481 bp deletion in the LRR domain (Liu et al., 2020). *bam1d-2* is a CRISPR-derived single base pair deletion in the kinase domain*. bam1d-3* is a Uniform Mu (UfMu) transposon insertion in the LRR domain. bp, base pairs, where numbers indicate bp position of different mutant alleles. (B) *bam1d-1* roots are insensitive to CLE7 and FCP1 peptide treatments. Plants were treated with 10 µM CLE7, FCP1, or a scrambled CLE7 peptide control (ScrCLE7). Letter codes indicate a statistically significant difference of p<0.05 using one way ANOVA with a post hoc Sidak test, n≥6. Experiment was repeated twice with similar results. (C) *bam1d-1* ears were shorter than WT. Scale bar, 2 cm. (D) Quantification of ear traits; *bam1d*-*1* ears were shorter with less kernels per row and lower ear and seed weight. Student t-test used to evaluate significance, * ≤; 0.05, ** ≤; 0.01, *** ≤; 0.001, n≥30. KNR, kernel number per row, KRN, kernel row number.

To determine whether *BAM1D* affects maize ear development, we assayed three *bam1d* mutant alleles (Figure 4A) derived from either genome editing or transposon insertion and found both *bam1d*-*1* and *bam1d*-*3* mutant ears were shorter than their WT siblings (Figure 4C, D Figure S7A). *bam1d-2* ears were not significantly shorter (Figure S7C), however this may be a weak allele because it has a lesion in the kinase domain, similar to weak *clv1* mutants (Diévart et al., 2003). *bam1d* mutants also had smaller inflorescence meristems (Figure 5A, B, Figure S8A, C). Together, these data show that *BAM1D* positively regulates ear and meristem size.

**Figure 5.**
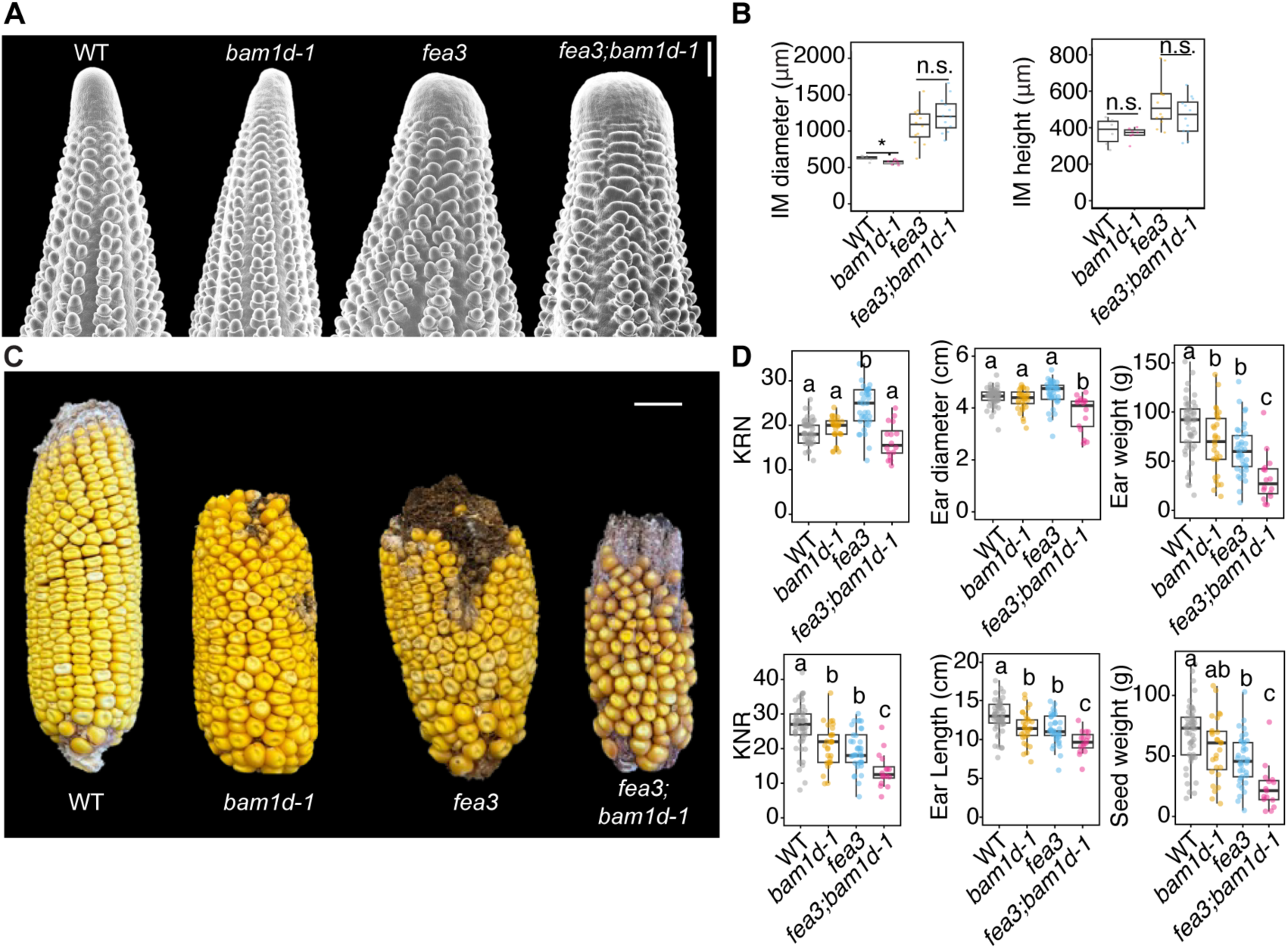
Complex epistatic and additive interactions between *fea3* and *bam1d*. (A) Scanning electron micrograph images of WT, *bam1d*-*1*, *fea3*, and *fea3*;*bam1d*-*1* ear primordia. (B) Inflorescence diameter and height measurements of WT, *bam1d*-*1, fea3,* and *fea3;bam1d* meristems. *bam1d-1* inflorescence meristems were narrower than WT, whereas *fea3;bam1d-1* meristems were not significantly different to *fea3*. p-values calculated using a Student’s t-test, n≥6. * indicates p<0.05, n.s. not significant. (C) *fea3;bam1d*-*1* ears are smaller than single mutant ears. Scale bar, 2 cm. (D) Quantification of ear traits in a segregating *fea3*;*bam1d*-*1* population. Kernel row number and ear diameter were decreased in *fea3;bam1d* compared to *fea3*, indicating epistasis for these two traits. In contrast, *fea3;bam1d-1* mutants had additive decreases in kernel number per row, ear length, ear weight, and seed weight. Letter codes indicate a statistically significant difference of p<0.05 using one-way ANOVA with post hoc Sidak test, n≥16. KNR, kernel number per row, KRN, kernel row number.

### bam1d and fea3 interact genetically to control ear development

Since FEA3 and BAM1D physically interact, we hypothesized that *fea3* and *bam1d* mutants would have a non-additive relationship if they operate in the same signal transduction pathway. We first assayed inflorescence meristem size in WT, *fea3*, *bam1d*, and *fea3;bam1d* plants. *bam1d* IMs were narrower than WT, however, IM diameter was not significantly different between *fea3;bam1d* and *fea3*, indicating that *fea3* is epistatic to *bam1d* in the control of IM diameter (Figure 5A, B, Figure S8B, C). We next observed mature ear phenotypes, and found epistasis for KRN and ear diameter traits. For example, *fea3;bam1d-1* mutants had a decrease in kernel row number (KRN) compared to *fea3,* even though *bam1d-1* had no effect on KRN (Figure 5C, D). *fea3*;*bam1d*-*2* and *fea3;bam1d-3* mutants also had a reduced KRN compared to *fea3* single mutants (Figure S7). However, some other traits were additive, such as kernel number per row, ear length, ear weight, and seed weight (Figure 5C, D). Overall, our results demonstrate non- additive and epistatic interactions between *fea3* and *bam1d,* suggesting they function in the same pathway in maize ear development. Intriguingly, the epistatic relationships between *FEA3* and *BAM1D* are different in the control of inflorescence meristem size and ear development, indicating FEA3 and BAM1D may be part of different signaling networks at different stages of inflorescence development.

## Discussion

FEA3 is a receptor-like protein required for meristem organization that acts in parallel with canonical CLV receptors CLV1/TD1 and CLV2/FEA2 (Je et al., 2016). Here we identified the receptor-like kinase BAM1D as a co-receptor of FEA3. BAM1D binds the same CLE peptide that FEA3 perceives; co-localizes with FEA3 in ear primordia; and physically interacts with FEA3 in shoot apices and ear primordia. *bam1d* mutants make shorter ears and have smaller inflorescence meristems. This echoes the reduced number of carpels and stamens observed in Arabidopsis higher order *bam* mutants, which results from a smaller meristem (DeYoung et al., 2006). The effect on ear size is modest in *bam1d* mutants, however, which may be a result of compensation from other *BAM* genes. Indeed, BAM1D-Turbo biotinylates BAM1B and BAM1C, and these receptors may compensate for a lack of BAM1D.

*bam1d* has a complex epistatic relationship with *fea3*, as some double mutant phenotypes are epistatic and others are additive. While *bam1d* has narrower IMs than WT, *fea3;bam1d* IM size is the same as *fea3,* demonstrating an epistatic effect on IM size. Epistasis is also observed in some mature ear traits, as KRN is reduced in *fea3;bam1d* compared to *fea3*. Furthermore, seed set is greatly reduced in *fea3;bam1d- 1* relative to *fea3*, which parallels Arabidopsis *bam* fertility defects (DeYoung et al., 2006; Hord et al., 2006; Nimchuk et al., 2015). While floral phenotypes can be interpreted as a direct readout of meristem activity, *CLV* genes have distinct roles in meristem regulation and floral development (Durbak and Tax, 2011; John et al., 2023). The epistatic interactions between *fea3* and *bam1d* in inflorescence meristem size and in the control of KRN are different, suggesting that *FEA3* and *BAM1D* may have distinct roles at different developmental stages. Because *BAM1D* is a QTL for seed size (Yang et al., 2019), and weak *fea3* alleles can increase KRN without a compensatory loss in seed size (Je et al., 2016), this further suggests unique roles for these genes in the control of inflorescence meristem architecture and seed development.

What molecular mechanism might explain the antagonistic *fea3* and *bam1d* phenotypes? One possibility is that FEA3 regulates the activity of BAM1D by increasing its binding affinity to FCP1 relative to other CLE peptides. This type of regulation is observed with the receptor-like protein TOO MANY MOUTHS (TMM) and receptor kinases ERECTA (ER) and ERECTA-LIKE 1 (ERL1) (Lin et al., 2017). TMM modifies the binding interface of ER and ERL, restricting the binding of certain EPFL peptides, but promoting the binding of others (Abrash et al., 2011). While TMM affects peptide binding, it does not directly bind EPFL peptides, similar to FEA3 interacting with BAM1D but not directly binding FCP1.

Alternatively, because BAM1D can perceive CLE7 in addition to FCP1, it may function in additional signaling pathways independently of FEA3. This is consistent with the observation that Arabidopsis *BAM* genes are pleiotropic developmental regulators, functioning in shoot, root, and vascular development, and bind to a wide range of CLE peptides (DeYoung et al., 2006; Guo et al., 2010). To further support the idea that BAM1D and FEA3 activity only partially overlaps, our proximity labeling interactomes did not fully overlap between FEA3 and BAM1D. FEA3 and BAM1D are also only partially co-localized in ear primordia. This suggests that while FEA3 and BAM1D form a receptor complex, they may function in additional signaling complexes. Signaling complex heterogeneity is observed with other CLV receptors, such as with FEA2, which interacts with different downstream signaling components in the perception of different CLE peptides (Je et al., 2018). Arabidopsis CLV2 and CRN can interact with BAMs, and CLV2/CRN promote the stability of BAM3 in the root (Hazak et al., 2017). Interestingly, Arabidopsis *clv2 bam* genetic interactions are similar to maize *fea3;bam1d* interactions, where carpel number is slightly repressed in *clv2 bam* mutants relative to *clv2* (DeYoung and Clark, 2008). Therefore, it is possible that BAM1D also interacts with FEA2 in an independent pathway from the FEA3-BAM1D receptor module (Figure 6B).

**Figure 6.**
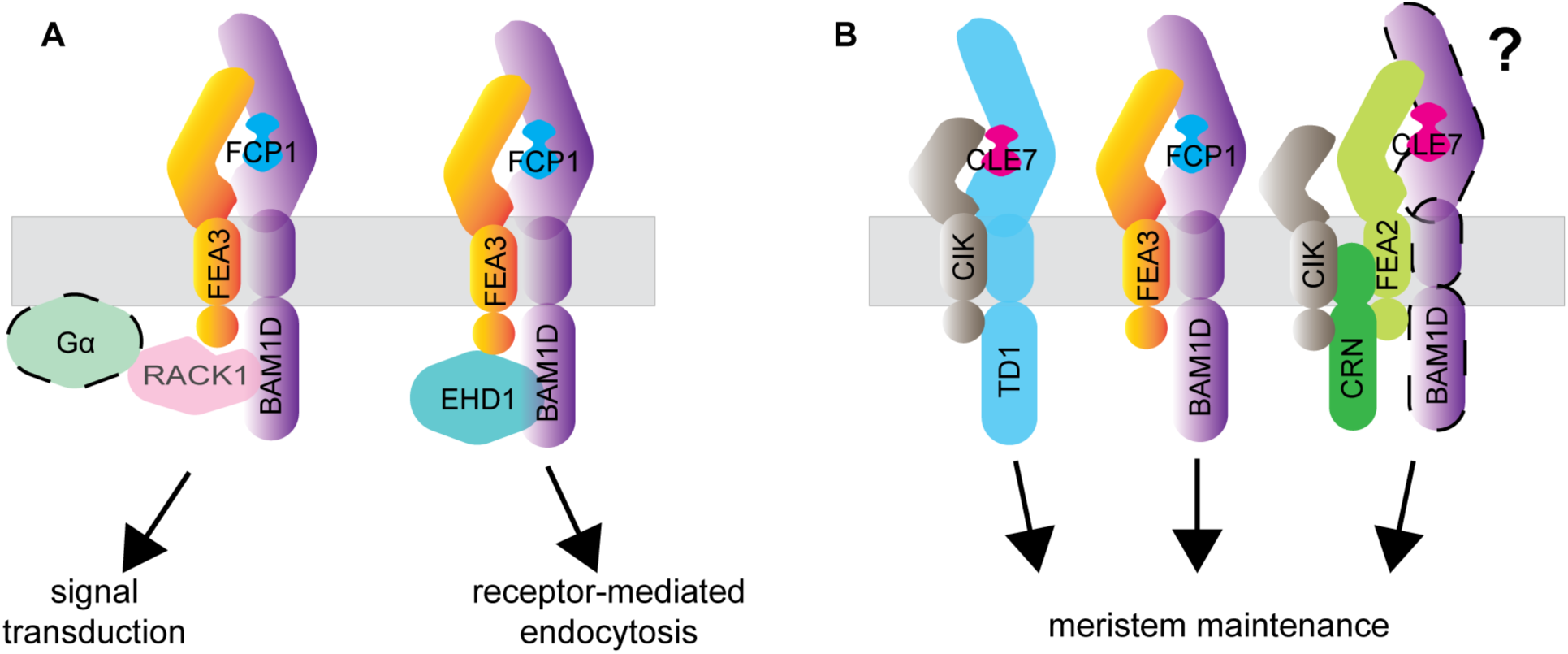
Proposed model for FEA3-BAM1D receptor signaling complex. (A) BAM1D binds FCP1. Binding affinity may be altered by FEA3, which allows signal transduction through G-protein signaling, since the G-protein scaffold RACK1 interacts with both FEA3 and BAM1D. FEA3 and BAM1D activity may be regulated by receptor-mediated endocytosis, as evidenced by internalized RFP-FEA3 and BAM1D-YFP signal, in addition to endocytic components such as EHD1 being labeled by FEA3 and BAM1D. (B) Meristem maintenance is mediated by several partially overlapping receptor complexes, which allows a quantitative readout of ligand perception.

TurboID AP-MS of FEA3-Turbo and BAM1D-Turbo revealed putative downstream signaling components of these receptor proteins, including several signaling-related proteins. This includes RACK1, a G-protein scaffold. G proteins have been implicated in CLV signaling, so this interaction suggests that FEA3 and BAM1D signaling is also mediated through heterotrimeric G-proteins (Bommert et al., 2013; Wu et al., 2020). Several proteins related to clathrin-mediated endocytosis were also enriched in both FEA3-Turbo and BAM1D-Turbo proximity labeling samples. It was recently found that a TPLATE component, AtEH1, interacts with CLV1 in Arabidopsis to attenuate CLV signaling through receptor-mediated endocytosis, so this suggests clathrin-mediated endocytosis is a widespread mechanism for regulating CLV signaling (Wang et al., 2023). Many proteins related to auxin signaling were also shared in both TurboID datasets, supporting connections between CLV signaling and auxin, as in Arabidopsis (Jones et al., 2021; John et al., 2023).

In summary, we propose that FEA3 and BAM1D interact to control maize ear size. Through proximity labeling, we show FEA3 and BAM1D interact *in vivo*, and identify several putative downstream signaling effectors of this signaling module. The partially overlapping proximity labeling targets of FEA3 and BAM1D and their antagonistic genetic interaction suggest they each function in multiple receptor complexes. Multiple BAM receptor complexes may help provide a quantitative readout of ligand perception to fine tune stem cell proliferation, since they can perceive a broad range of CLE peptides (Figure 6B). This receptor signaling paradigm appears broadly conserved, as several mammalian receptors operate in heterogenous complexes (Su et al., 2022; Granados et al., 2024). Future work should probe how BAM receptor complexes vary in different tissue types, to clarify receptor complex heterogeneity. Given the dazzling array of LRR receptors in plants (Man et al., 2020), it is clear we have only begun to scratch the surface of receptor-mediated control of development.

## Methods

### Plant growth

Maize plants were grown at Cold Spring Harbor, NY (40° 51′ N 73° 27′ W), Lloyd Harbor, NY (40° 54′ N 73° 28′ W), or Riverhead, NY (40° 57’ N 72° 43’ W) from June to October for phenotyping in summer of 2023 and 2024. Material for TurboID experiments was grown in the greenhouse under long-day conditions (16 h light, 8 h dark) at 28° C.

### Generation of mutants, genotyping, and phenotyping

*fea3-0* is a partial transposon insertion allele previously published in Je et al., 2016. *Zmbam1d-1* is a 458 bp deletion in the first exon generated by CRISPR previously published in Liu et al., 2020. *Zmbam1d-2* is a CRISPR allele generated in the B104 background. CRISPR guides were designed using CRISPR-P (Liu et al., 2017) and synthesized in an array into pDONR using Gene Universal, with the sgRNA sequence GACGTGTACAGCTTCGGCG used to target *BAM1D*. Arrays were cloned into backbone pMCG1005 using Gateway cloning. *Zmbam1d-3* is a UFMu insertion from the W22 background. All *bam1d* alleles were introgressed into *fea3*-*0* (B73 background) for at least two generations before phenotyping.

To confirm the presence of *fea3-0*, we used PCR with primers PL108 and PL109 to amplify the insertion. For *bam1d-1*, we used primers PL29 and PL30. To genotype *bam1d-2*, we used a dCAPS marker assay, amplifying a PCR product with PL25 and PL26, and then performing a restriction digest with Hpy188I to cut the WT allele. To genotype *bam1d-3*, we used primers PL27 and PL28 to amplify the WT sequence, and PL168 (TIR6) and PL27 to amplify the Mu insertion. All primer sequences are listed in Supplementary Table 4.

Segregating populations were used for phenotyping to ensure a uniform background. All statistical analyses were performed in RStudio using R version 4.4.0.

### Cloning

We used Gateway cloning to generate all translational fusions in this study, listed in Supplementary Table 5. To generate BAM1D:BAM1D-YFP, BAM1D:BAM1D-Turbo, and FEA3pr:FEA3-Turbo constructs, we used four-way Gateway cloning. We cloned promoters into p1p5, genomic sequences into p5p4, YFP or Turbo into p4p3, and 3’UTRs into p3p2. The TurboID coding sequence was codon optimized for maize. We added a flexible linker to the N terminus of the coding sequence, and a C-terminal FLAG tag to be able to detect the protein via Western blot. The coding sequence was synthesized (Gene Universal) and subcloned into Gateway pDONR p4p3 to generate c-terminal translation fusions with FEA3 and BAM1D. The primers used to generate attb fragments for cloning into pDONR vectors are listed in Supplementary Table 4. 35S:BAM1D-FLAG was cloned using pCAMBIA1300, and 35S:FEA3-MYC was generated using pAM1006 as the destination backbone. The FEA3 sequence was codon-optimized for *N. benthamiana* to improve expression.

### Photoaffinity labeling

To perform Halo binding assays, Halo translational fusions were generated for FEA3-Halo, AtBAM1-Halo, or BAM1D-Halo using Gateway cloning, primers are listed in Supplementary Table 4.

Photoactivatable [^125^I]ASA-FCP1 was prepared as follows. The Fmoc-protected FCP1 analog Fmoc-[Lys^2^]FCP1 was synthesized by Fmoc chemistry using a peptide synthesizer. [(4-azidosalicyl)Lys^2^]FCP1 (ASA-FCP1) was prepared by coupling 4- azidosalicylic acid succinimidyl ester and Fmoc-[Lys^2^]FCP1 as described (Ogawa et al., 2008). ASA-FCP1 was further radioiodinated by the chloramine T method. Labeled peptide was purified by reverse-phase HPLC to yield analytically pure [^125^I]ASA-FCP1 with specific radioactivity of 28 Ci/mmol.

ZmBAM1D-Halo and FEA3-Halo were expressed in tobacco BY-2 cells as described (Ogawa et al., 2008). Photoaffinity labeling, immunoprecipitation, SDS-PAGE and autoradiography was performed according to previous report (Shinohara et al., 2016) using 90 nM [^125^I]ASA-FCP1. For the competitive binding assay, 300-fold molar excesses of unlabeled peptides were used.

### Peptide treatment assay

To perform CLE peptide treatment assays, mature seeds were surface sterilized and grown on MS agar for 2 days in the dark to allow germination. Seedlings with similar root emergence were selected and transferred to agar media containing 10 µM CLE peptide, sequences listed in Supplementary Table 6. Seedlings were grown on peptide media for 7 days, after which they were scanned, and root length measurements were performed in Fiji using the Measure tool.

### in situ hybridization

*ZmBAM* expression in the developing ear was characterized by mRNA *in situ* hybridization assays. Developing ears (5mm) of B73 were collected and fixed for paraffin embedding. The mRNA probes used for mRNA *in situ* hybridization were amplified by *ZmBAM* gene-specific primers listed in Supplementary Table 4. A previously described protocol was employed to detect the expression of *ZmBAMs* (Jackson et al., 1994).

### TurboID

For the TurboID AP-MS experiment, shoot apices with surrounding immature leaves were dissected on ice from FEA3-Turbo, BAM1D-Turbo, and non-transgenic plants 2 weeks post planting. Five hundred mg of tissue was treated with 50 µM biotin for 2 h at room temperature. Following biotin treatment, tissue was rinsed three times with ice cold water and frozen at -80 °C.

For protein extraction, 1 ml of cold protein extraction buffer (50 mM Tris, pH 7.5, 150 mM NaCl, 0.1% SDS, 1% Triton-X 100, 0.5% Na-deoxycholate, 1 mM EGTA, 1 mM DTT, 1x cOmplete protease cocktail inhibitor, 1 mM PMSF) was added to 500 mg of frozen homogenized tissue in a pre-chilled mortar and pestle. Tissue was homogenized with the extraction buffer using the mortar and pestle, allowed to thaw, and transferred to Eppendorf tubes. Tubes were rotated on a rotor wheel for 10 min at 4 °C. Samples were sonicated in the cold room using a Branson SFX150 Sonifier at 30% power, with 0.5 sec pulses for 20 sec. Samples were then centrifuged for 5 min at 1,500 rpm at 4 °C. The supernatant was transferred to a new tube and centrifuged at 12,000 rpm for 10 min at 4 °C. The supernatant was transferred to a new tube. Following protein extraction, free biotin was removed using PD10 desalting columns. Samples were eluted, and protein concentration was measured using a Pierce assay (Thermo Scientific).

Biotinylated proteins were purified from the protein lysate using 200 µl MyOne Streptavidin C1 (Invitrogen) magnetic beads using an overnight incubation at 4 °C. To wash the beads, beads were spun down briefly at 1500 rpm. Beads were washed twice with extraction buffer, twice with equilibration buffer (50 mM Tris pH 7.5, 150 mM NaCl, 0.1 % SDS, 1 % Triton-X-100, 0.5 % Na-deoxycholate, 1 mM EGTA,1 mM DTT), once with cold 1 M KCl, once with cold 100 mM Na2CO3, and once with 2 M Urea in 10 mM Tris pH 8.0 at room temperature, with eight minute incubations on the rotator for each wash. Following the washes, beads were resuspended in extraction buffer and sent for tryptic digest.

For on-bead tryptic digestion, the streptavidin beads were washed seven times with PBS (8 minutes each wash), then incubated for 3 hours at 25°C in trypsin buffer (50 mM Tris pH 7.5, 1 M urea, 1m M DTT, 0.4 µg Trypsin). The supernatant from this initial digestion, along with two subsequent 60 µL washes with 1 M urea in 50 mM Tris pH 7.5, was pooled. The combined eluates were reduced, alkylated and digested overnight with 0.5 µg trypsin. An additional 0.5 µg of trypsin was added in the next morning, followed by acidification with formic acid 4 hours later to a final concentration of ∼ 1 %. Peptides were desalted using OMIX C18 pipette tips.

LC-MS/MS was carried out on a Orbitrap Eclipse Tribrid mass spectrometer (Thermo Fisher), equipped with an Easy LC 1200 UPLC liquid chromatography system (Thermo Fisher). Peptides were first trapped using a trapping column (Acclaim PepMap 100 C18 HPLC, 75 μm particle size, 2 cm bed length), then separated using analytical column AUR3-25075C18, 25CM Aurora Series Emitter Column (25 cm x 75 µm, 1.7 µm C18) (IonOpticks). The flow rate was 300 nL/min, and a 120-min gradient was used. Peptides were eluted by a gradient from 3 to 28 % solvent B (80 % acetonitrile, 0.1 % formic acid) over 106 min and from 28 to 44 % solvent B over 15 min, followed by a short wash (9 min) at 90 % solvent B. Precursor scan was from mass-to-charge ratio (m/z) 375 to 1600 (resolution 120,000; AGC 200,000, maximum injection time 50ms, Normalized AGC target 50%, RF lens(%) 30) and the most intense multiply charged precursors were selected for fragmentation (resolution 15,000, AGC 5E4, maximum injection time 22ms, isolation window 1.4 m/z, normalized AGC target 100%, include charge state=2-8, cycle time 3 s). Peptides were fragmented with higher-energy collision dissociation (HCD) with normalized collision energy (NCE) 27. Dynamic exclusion was enabled for 30s.

For LFQ analysis, a MaxQuant (ver. 1.6.2.10) search was executed using default parameters with the following changes: In Group-specific parameters Label-free quantification was enabled with “Fast LFQ” checked. In global parameters “Match between runs” were enabled. For Identification, peptides were searched against the Zm- B73-REFERENCE-GRAMENE-4.0 protein database (Downloaded from maizegdb.org 05/03/19) containing a total of 143,679 entries. The proteingroups.txt file output from MaxQuant was analyzed in Perseus (ver. 2.0.10.0). LFQ intensities were imported and filtered with the following features: removing ‘reverse = +’, ‘potential contaminant = +’ and ‘only identified by site = +’. Data was log2 transformed and rows that were not identified in at least three replicates of one sample group were removed. Missing values were imputed from normal distribution (width = 0.3, down shift = 1.8, total matrix mode). A Student’s t-test was executed to examine protein groups with different intensities between the elute and control groups. The t-test settings were the following: ‘Permutation-based FDR’, ‘FDR=0.05’,S0=1, ‘Report q-value’, ‘Number of Randomizations=250’ and ‘-log10 p-value’.

Proteins with a log2fc of 1 and q-value ≤; 0.05 were considered statistically significant. GO enrichment was assessed using ShinyGO 0.81 (https://bioinformatics.sdstate.edu/go/). Volcano plots and bubble plots were generated in RStudio using ggplot2. Venn diagrams were generated with the eulerr package in RStudio. Hypergeometric scores were calculated using the phyper function in R.

For the targeted TurboID experiment, 5-10 mm ear primordia were dissected from maize plants, treated with 250 µM biotin for 1 h at room temperature, rinsed three times with ddH2O, and frozen at -80 °C. For protein extraction, samples were homogenized in a pre-chilled mortar and pestle, and two volumes of extraction buffer was added to 1 volume of tissue. Protein extraction was performed as described for the TurboID AP-MS, except the desalting step was not performed. Following protein extraction, 500 µl of the protein extract was incubated with 20 µl GFP magnetic resin with end-over-end shaking for 1 h (Chromotek). Following immunoprecipitation, samples were washed three times with wash buffer (50 M Tris-HCl pH 7.5, 150 mM NaCl, 1% Triton-X, 1 mM EGTA). Beads were transferred to a new tube following the third wash, and immunoprecipitated proteins were eluted by incubating beads in 80 µl 2X Laemmli buffer with 2.5% beta-mercaptoethanol (BME) at 95 °C for 10 min. 2% of the input fraction and 25% of the immunoprecipitated fraction were loaded on the SDS-PAGE gel for immunoblot analysis.

For the biotinylation time-course, either five 5 mm ear primordia or five 2 week old shoot apices were pooled, treated with either 100 or 250 µM biotin, rinsed, homogenized in liquid nitrogen, and resuspended in two volumes of RIPA extraction buffer. After a 15 min incubation on ice, samples were centrifuged for 10 min at 12,000 g. 1 volume of 2X Laemmli buffer plus 2.5% BME was added to the supernatant. Samples were boiled for 10 min at 95 °C, spun down for 2 min at 10,000 g. 10 µg of protein was loaded per sample. For immunoblot analysis, protein samples were loaded on 10 % SDS-PAGE gels.

Gels were run for 2 h at 100 V. For immunoblotting, PVDF membranes (Immobilon) were activated for 5 minutes in methanol. Proteins were transferred for 2 h at 100 V at 4 C using a BioRad transfer cell. Blots were blocked for 1 h in either 3% milk or 3% BSA on a tilting shaker. Immunoblots were then incubated at 4 °C overnight in primary antibody solution (SA-HRP, 1:100,000 in 3% BSA-PBST, anti-FLAG-HRP, 1:2,000 in 3% milk-PBST, anti-GFP (Roche), 1:2,000 in PBST). Blots were then rinsed four times with PBST, ten minutes each at room temperature. Blots were next incubated with secondary antibody solution for 1 hour at room temperature, followed by four rinses with PBST. Chemiluminescent ECL substrate was added to blots and incubated for 5 min at room temperature in the dark. Westerns were imaged using a BioRad imager.

### Confocal microscopy

To image BAM1D-YFP and RFP-FEA3 translational fusions, we used a Zeiss LSM 900 confocal microscope. To image ear primordia, we dissected 2-5 mm ears, embedded ears in 6% agarose, and made 100 µm sections using a vibratome. Shoot apices were hand dissected under a stereoscope and then mounted on a microscope slide. To image maize roots, emerging lateral roots were selected and imaged. To limit endocytosis of BAM1D-YFP and RFP-FEA3, we treated samples with 2 µM D15 peptide for 2 h as performed in Je et al., 2016. To image RFP-containing constructs, fluorophores were excited with a 561 nm laser, and emission spectra were collected from 565-700 nm with a GaAsP-Pmt2 detector. To image YFP constructs, fluorophores were excited with a 488 nm laser, and emission spectra were collected from 500 to 555 nm with a GaAsP-Pmt1 detector.

### Scanning electron microscopy

Two to five mm ear primordia were harvested from maize plants and attached to metal stubs with double-sided tape. SEM images were taken using a Jeol JCM-7000 benchtop scanning electron microscope with the following conditions: high vacuum, 15 kV, high probe current, analysis mode. Meristem measurements were made using Fiji.

### Gene tree inference

Primary transcript peptide annotation databases for *Arabidopsis thaliana*, *Amborella trichopoda*, *Brachypodium distachyon*, *Oryza sativa*, *Solanum lycopersicum*, *Setaria viridis, Sorghum bicolor* and *Zea mays* were obtained from Phytozome v.12 (Goodstein et al., 2012). LRR-RLK Clade XI search priors were collected from Dufayard *et al*. using genes classified in Clade XI from *A. thaliana*, rice, tomato, and *B. distachyon*. These search priors were aligned using the L-INS-i algorithm in Mafft v.7.313 (Katoh et al., 2013), then used to build a hidden Markov model (HMM) profile. The HMM profile was applied with Hmmer v.3.1b2 (Eddy et al., 2011) to search the peptide databases. Newly identified LRR-RLKs were re-aligned in MAFFT as previously and used to infer a maximum likelihood gene tree with IQ-tree v. 0.9. 3 (Nguyen et al., 2015) employing default ModelFinder settings and 1000 bootstrap replicates. The resulting tree was visualized using Figtree v1.4.4 in the BEAST package (Bouckaert et al., 2014).

### AlphaFold-Multimer predictions

AlphaFold-Multimer predictions were generated in ChimeraX-1.9 using the AlphaFold structure prediction tool (Meng et al., 2023; Abramson et al., 2024). The LRR domain of BAM1D and mature peptide sequence of FCP1 or a scrCLE control were used for the predictions.

## Author contributions

PLL wrote the manuscript and performed all experiments except as indicated below. FX performed the initial genotyping of *bam1d* mutants, and cloned constructs for the Halo experiment. LL performed the *in situ* hybridization experiments, and initial genotyping and characterization of *bam1d* mutants. PB performed root CLE peptide treatment experiments, TurboID time-course experiments, and assisted with piloting the maize TurboID AP-MS experiment and data analysis. BJ performed the *fea3;td1* genetic analysis. AR and SX performed the tryptic digest and mass-spectrometry analysis. JM and MB performed phylogenetic analysis of BAM genes. TS cloned BAM1D-YFP, and BAM1D-Turbo constructs. YM performed peptide binding assays. DJ edited the manuscript, supervised the project and provided funding for the project.

## Acknowledgements

Funding for this work was supported by USDA-NIFA Award No. 2020-67013-30909. PLL was supported by NSF Postdoctoral Research Fellowships in Biology Program under Grant No. 2010642. FX was supported by HFSP Long-Term fellowship program (LT000227/2016-L) and Natural Science Foundation of Shandong Province (ZR2023JQ012). BJ was supported by the National Research Foundation of Korea (NRF) grant funded by the Korea government (RS-2021-NR058971). This work was also supported by the National Institute of Health (S10OD030441 to S-L.X.) and by the Carnegie endowment fund for the Carnegie Mass Spectrometry Facility. Thank you to the Pedmale lab for use of their confocal microscope, and the Lippman lab for use of their scanning electron microscope. We thank Tim Mulligan, Kyle Schlecht, Autumn Harrison for their assistance with plant care. Many thanks to Jackson Lab undergraduate field interns for their support with DNA extractions and genotyping.

## Supplemental figures

**Supplemental figure 1.**
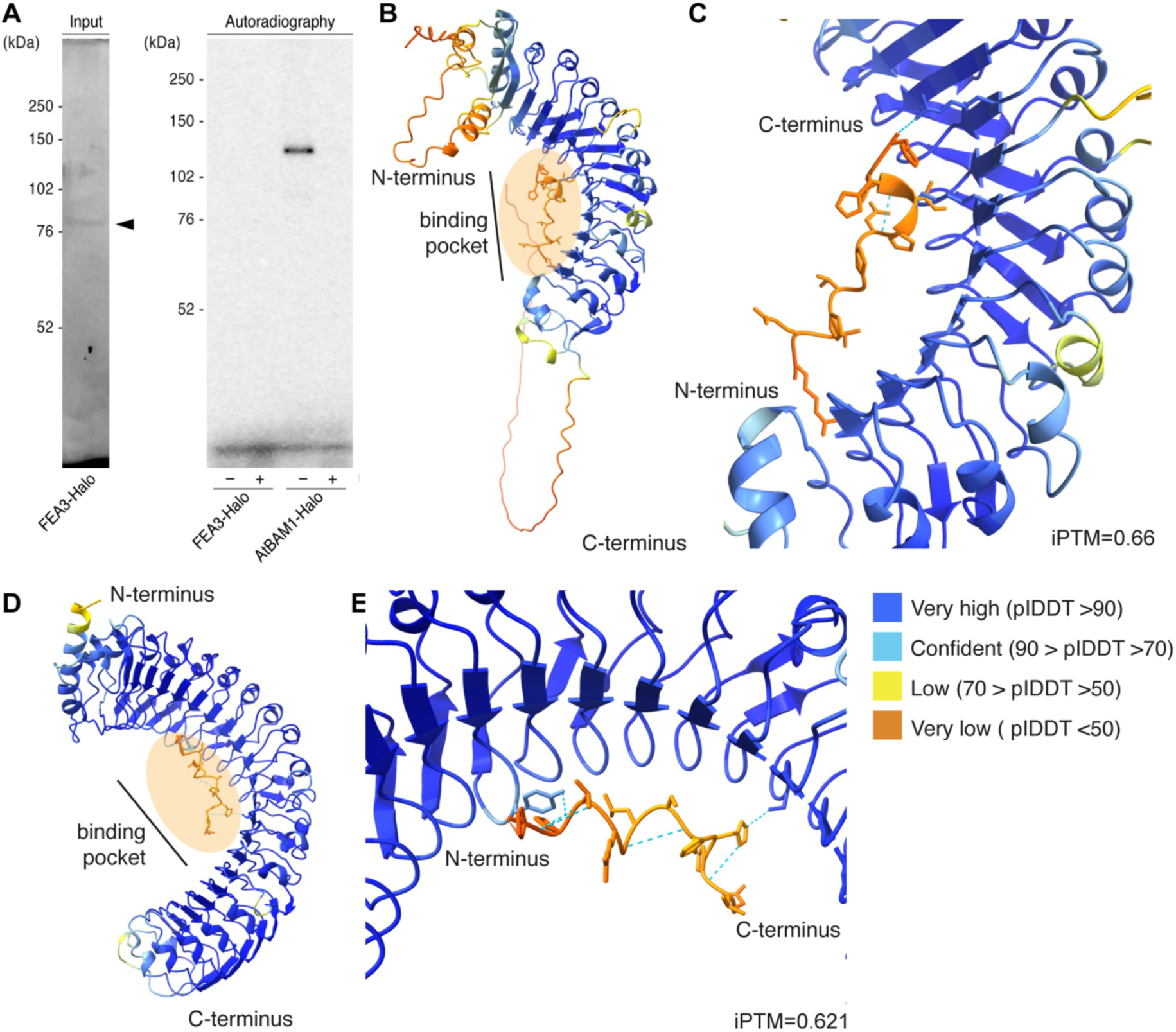
FEA3 does not bind FCP1. (A) Halo binding assay between FCP1 and FEA3-Halo or AtBAM1-Halo. FCP1 binds arabidopsis (At) BAM1, but not FEA3. – or + indicate absence or presence of unlabeled FCP1 peptide competitor. Arrow points to FEA3-Halo input band. (B) Overview of AlphaFold-Mutlimer predicted interacton between FEA3 extracellular domain and FCP1. FEA3 LRR and FCP1 are colored by Local Distance Difference Test (pIDDT) score, where bluer values indicate a stronger confidence prediction, and yellow and orange values indicate a weaker confidence prediction. (C) Close-up of binding interface between FEA3 and FCP1. Only one hydrogen bond is predicted between FCP1 and FEA3, represented by dashed lines. iPTM is a calculation of the predicted interaction likelihood between FCP1 and FEA3, with values above 0.8 indicating a strong confidence prediction. (D) Overview of predicted interacton between BAM1D and scrCLE. BAM1D and scrCLE are colored by Local Distance Difference Test (pIDDT) score, where bluer values indicate a stronger confidence prediction, and yellow and orange values indicate a weaker confidence prediction. (E) Close-up of binding interface between BAM1D and scrCLE. Only two hydrogen bonds are predicted between scrCLE and BAM1D, represented by dashed lines. iPTM is a calculation of the predicted interaction likelihood between scrCLE and BAM1D.

**Supplementary figure 2.**
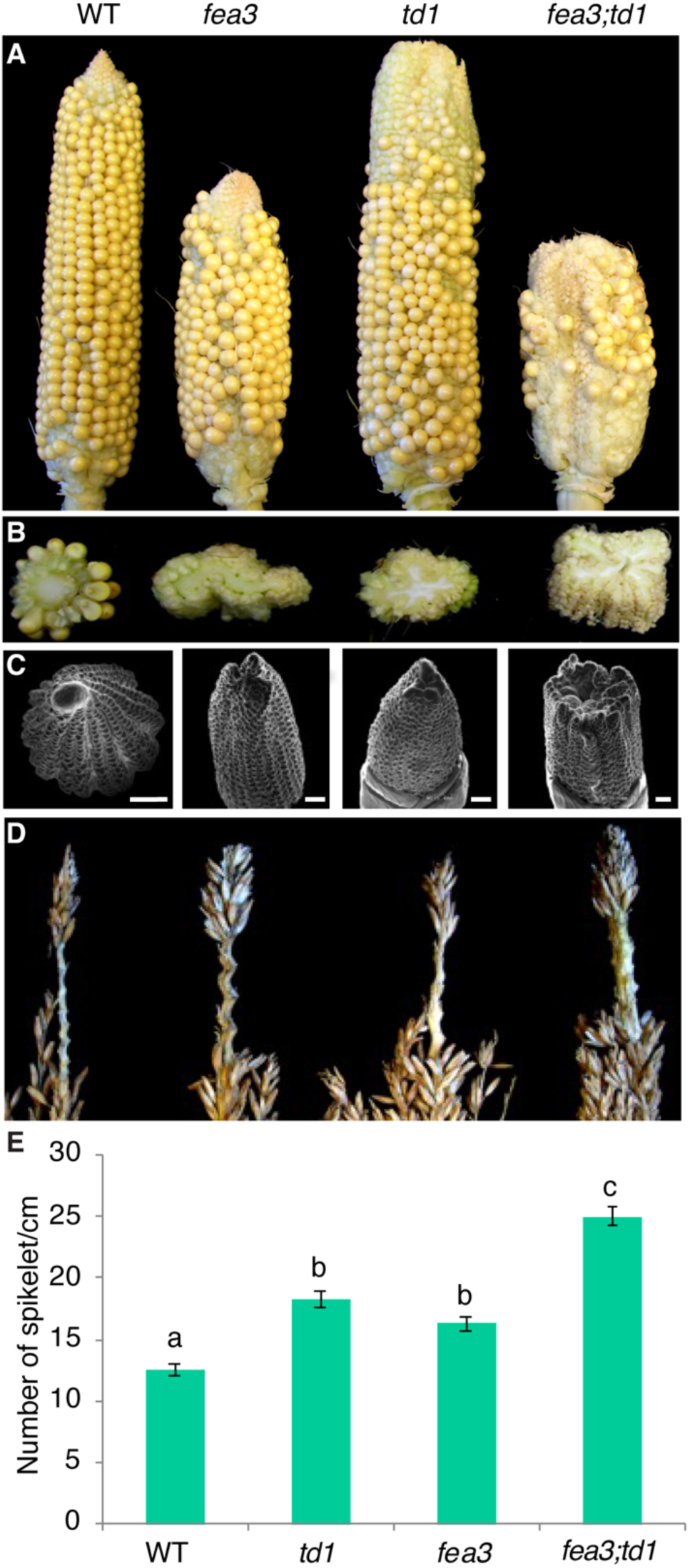
*fea3* and *td1* additively affect maize ear and tassel development. (A, B) *fea3;td1* ears were additively fasciated compared to *fea3* and *td1*. A, images of ears, B, cross-section through ears. (C) *fea3;td1* inflorescence meristems were additively larger compared to *fea3* and *td1*. Scale bar, 500 µm (D, E) *fea3;td1* spikelets were additively denser compared to *fea3* and *td1*. Letters indicate statistically significant difference, one-way ANOVA, n≥10.

**Supplemental figure 3.**
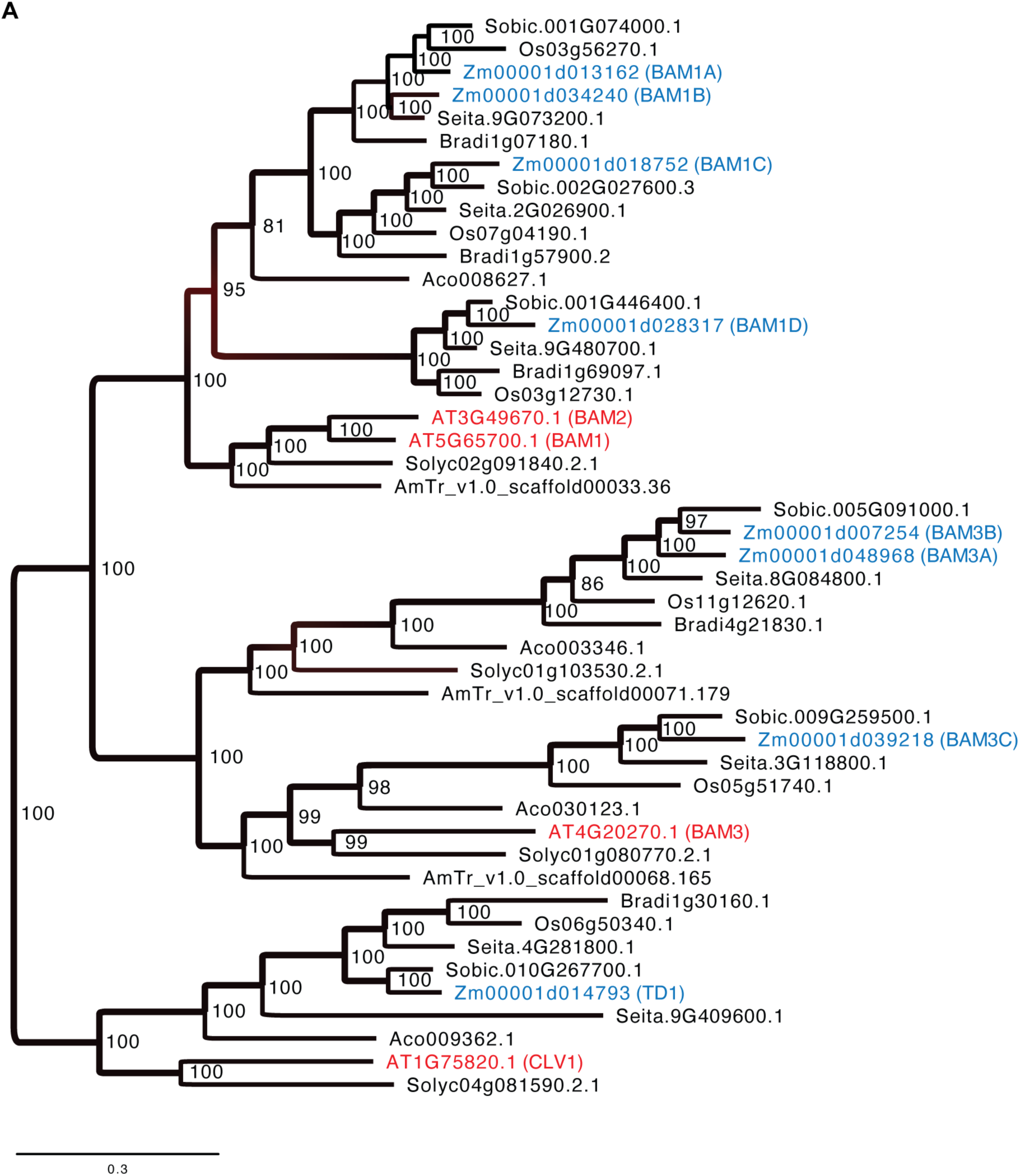
*BAM* gene tree. (A) Maximum likelihood *BAM* gene tree. Arabidopsis *BAM* orthologs are highlighted in red, maize *BAM* orthologs are highlighted in blue. Values on the tree indicate bootstrap support.

**Supplemental figure 4.**
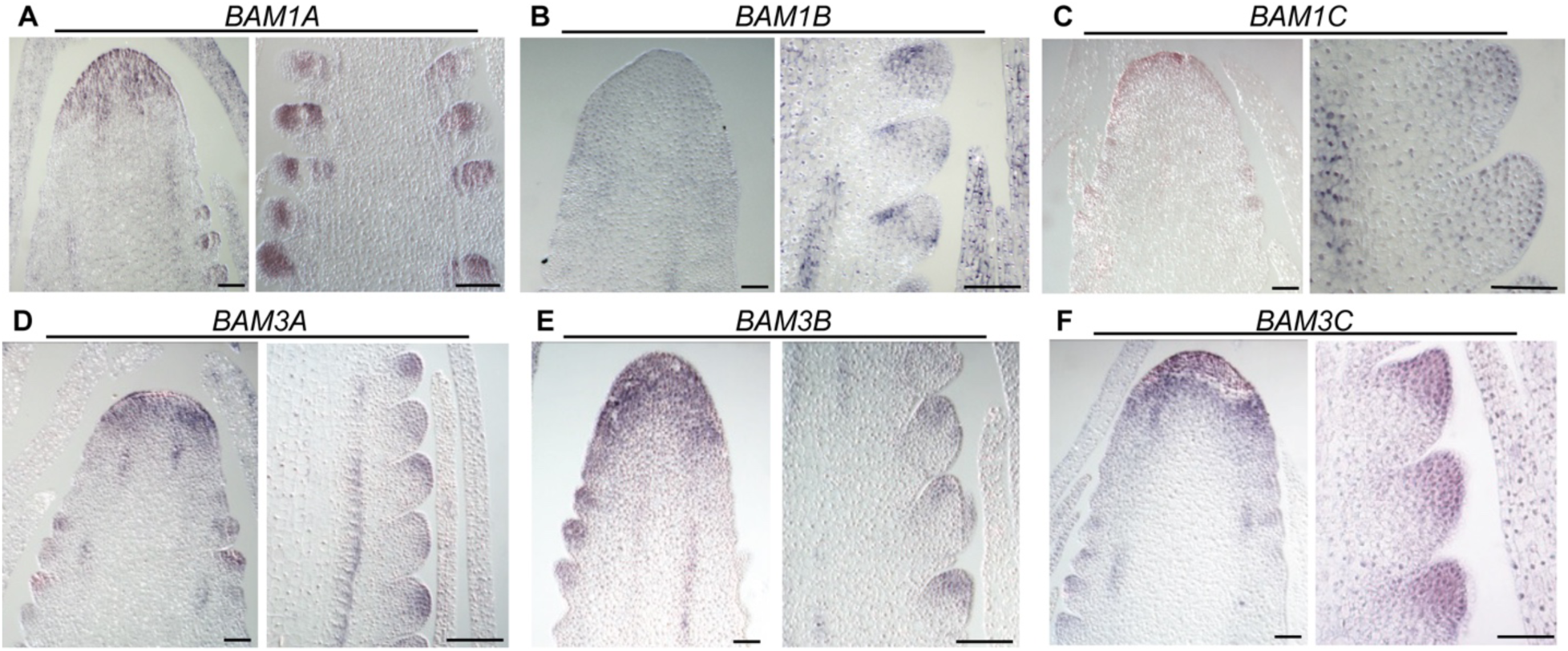
*in situ* hybridization of maize *BAM* genes in ear primordia. *BAM1A*, *BAM3A*, and *BAM3B* were broadly expressed in the inflorescence meristem and spikelet meristem. *BAM1B* was expressed mostly highly in the flanks of the spikelet meristem. *BAM1C* was expressed in the epidermis of the inflorescence meristem and spikelet meristem. Antisense probes. Scale bars are 50 µm.

**Supplemental figure 5.**
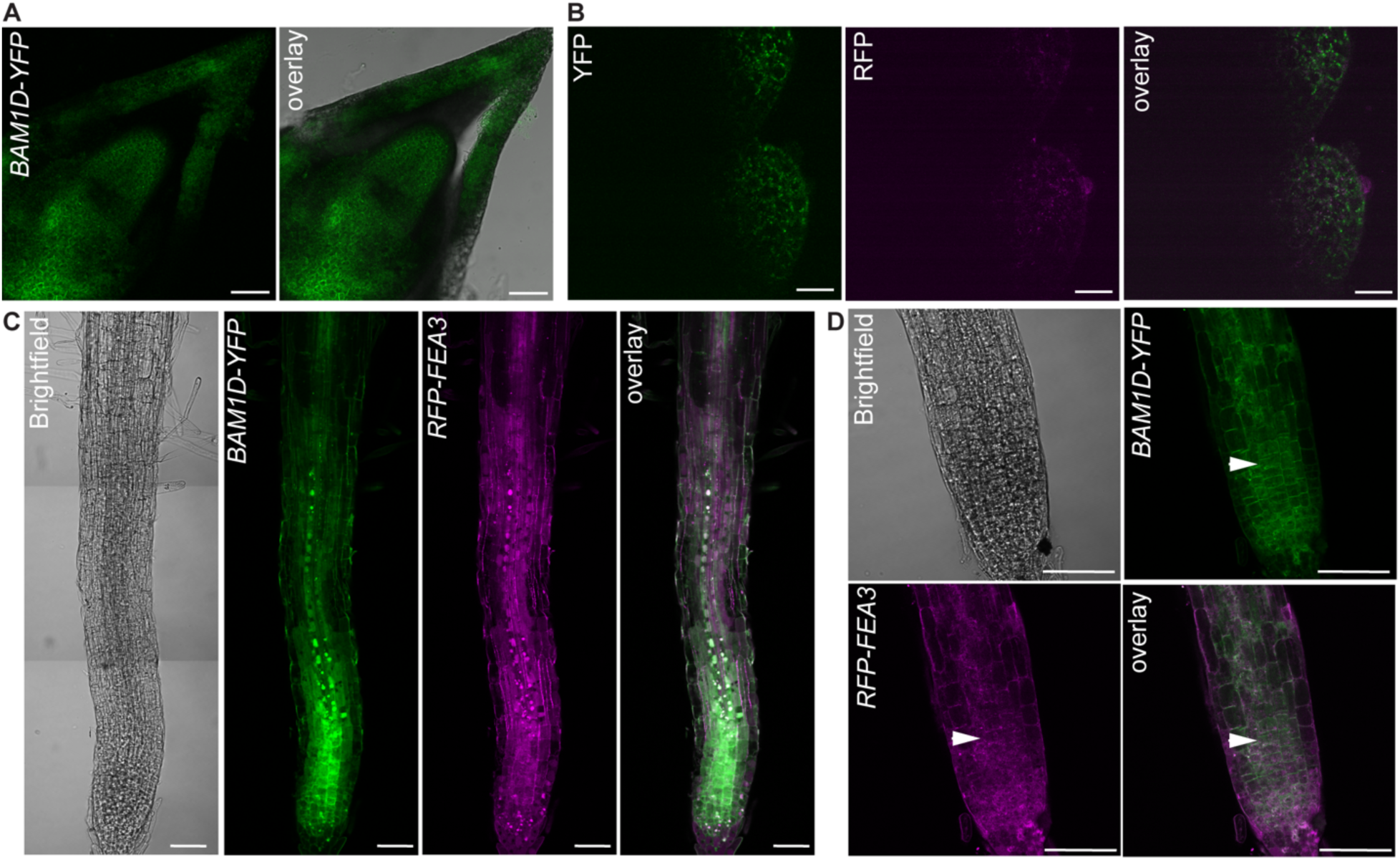
BAM1D-YFP is localized in inflorescence, shoot, and root meristems. (A) BAM1D-YFP was expressed broadly in the shoot apical meristem. (B) YFP and RFP autofluorescence in non-transgenic spikelet meristem. (C, D) BAM1D-YFP was expressed in the root apical meristem and partially co-localized with RFP-FEA3. Arrows indicate cell with co-localized signal. Scale bars are 50 µm.

**Supplemental figure 6.**
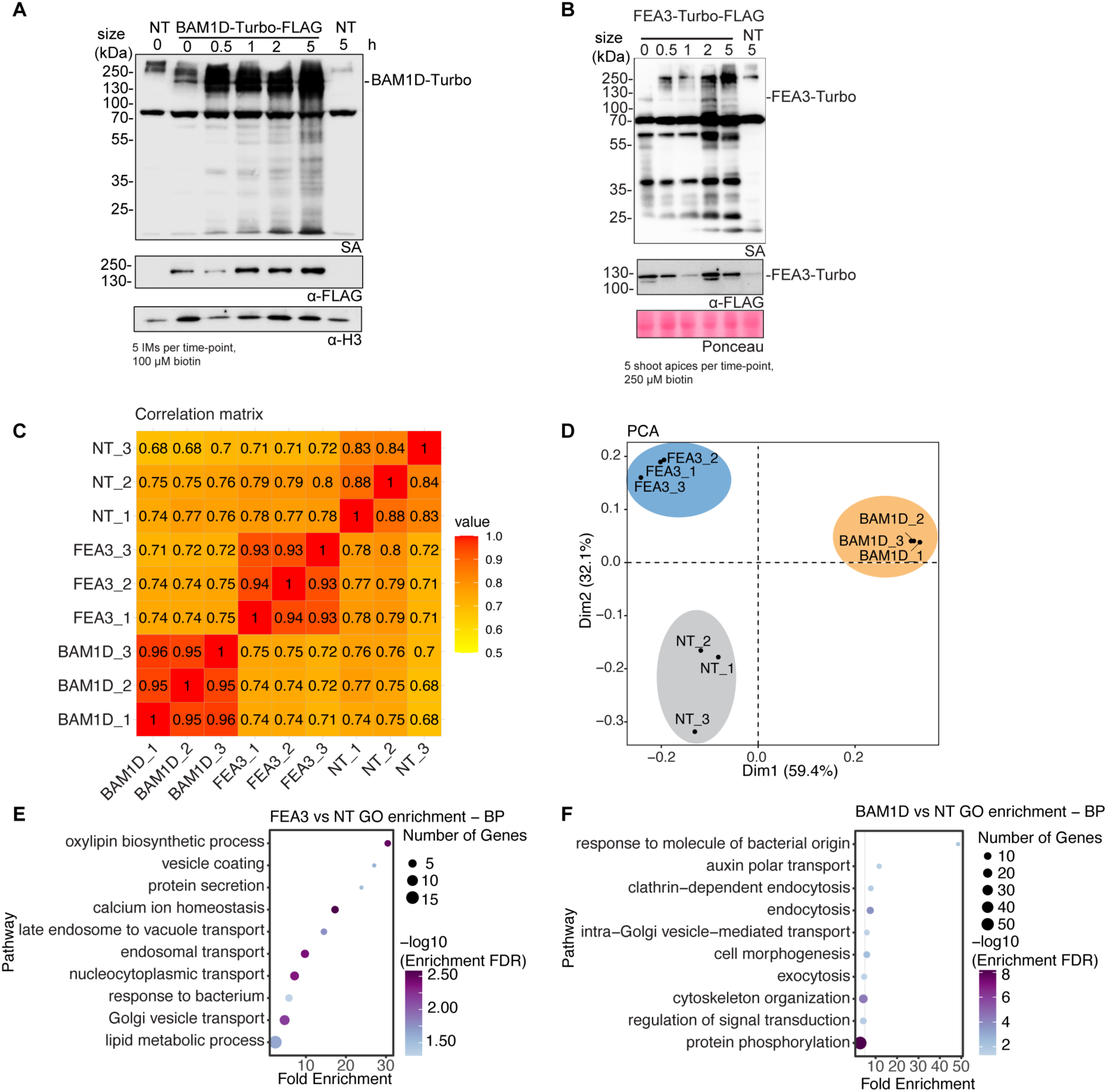
TurboID time-course and AP-MS. (A) Western blot showing time-course biotinylation experiment of BAM1D-Turbo in inflorescence meristem tissue. (B) Western blot showing time-course biotinylation experiment of FEA3-Turbo in shoot meristem tissue. NT, non-transgenic, h, hours, SA, streptavidin, H3, histone H3. (C) Correlation analysis of TurboID AP-MS samples. Numbers in matrix indicate the correlation strength between samples. (D) Principal component analysis showing the clustering of TurboID AP-MS biological replicates. (E) Significant GO term enrichment of FEA3-Turbo enriched proteins, BP, biological process, NT, non-transgenic. (F) Significant GO term enrichment of BAM1D-Turbo enriched proteins, BP, biological process, NT, non- transgenic.

**Supplemental figure 7.**
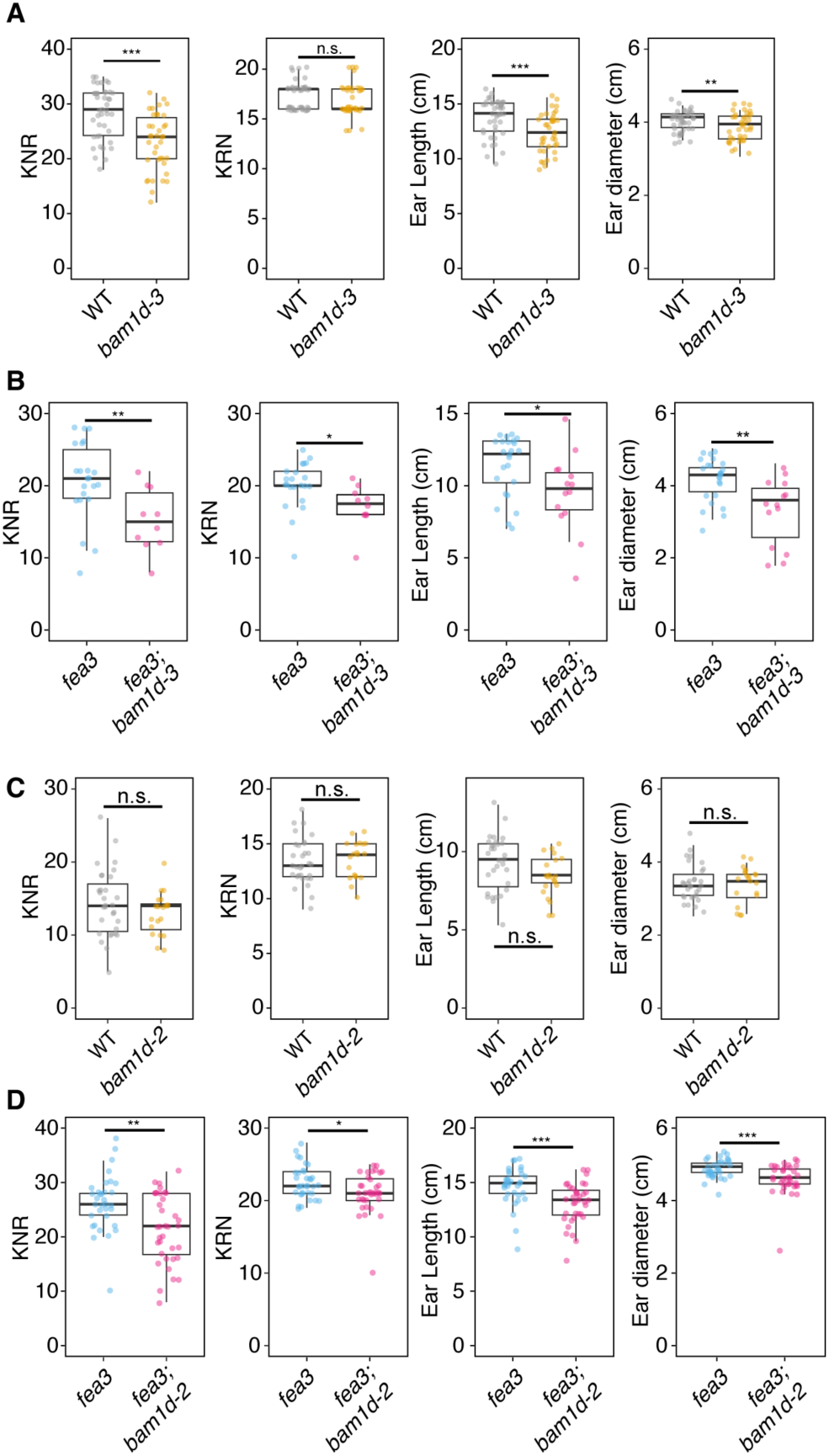
Ear size was not affected in *bam1d-2* and *bam1d-3* ears were smaller than WT. (A) Ear trait measurements in segregating *bam1d*-*3* populations. KNR, ear length, and ear weight were reduced in *bam1d*-*3* relative to WT. n=46. (B) Ear size was reduced in *fea3;bam1d-3* compared to *fea3*. n≥16. (C) Ear size was unaffected in *bam1d-2*. n≥21. (D) Ear size was reduced in *fea3;bam1d-2* compared to *fea3*. n≥34. * indicates p<0.05, ** indicates p<0.01, *** indicates p<0.001, n.s. not significant. Different letters indicate statistically significant difference p<0.05, Student t-test. KNR, kernel number per row, KRN, kernel row number.

**Supplemental Figure 8.**
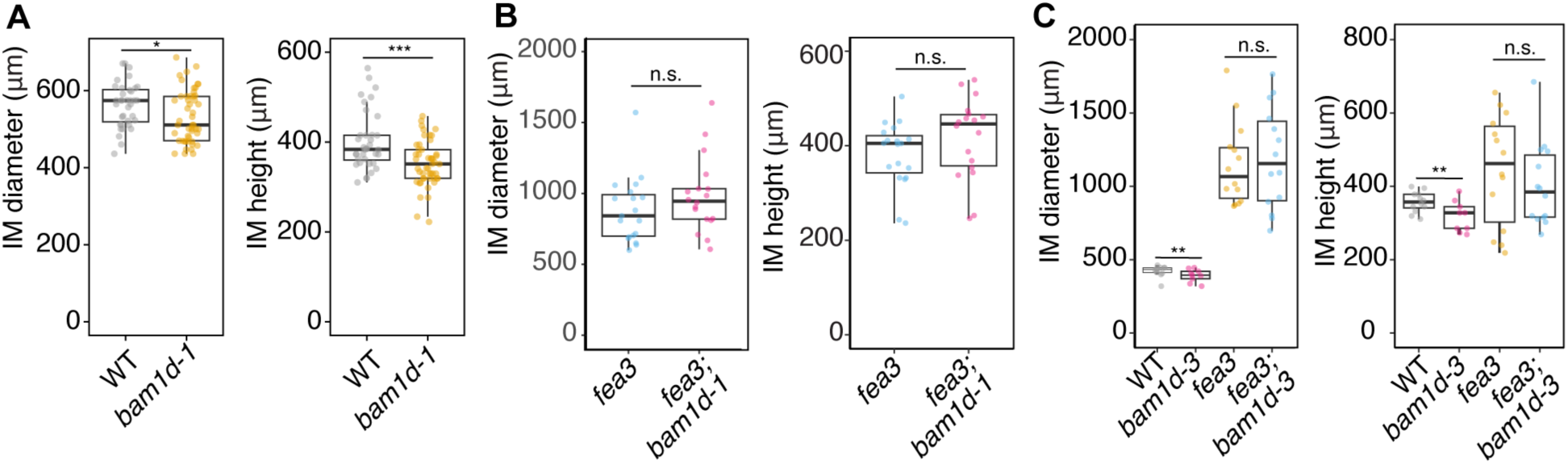
b*a*m1d mutants have smaller meristems, and *bam1d* is epistatic to *fea3* in the control of meristem size. (A) Inflorescence diameter and height measurements of WT and *bam1d*-*1* meristems. *bam1d-1* inflorescence meristems were shorter and narrower than WT. n≥36. (B) Meristem measurements of *fea3* and *fea3;bam1d-1* meristems. *fea3;bam1d-1* meristem size was not significantly different than *fea3*. n≥18. (C) Inflorescence diameter and height measurements of WT, *bam1d-3*, *fea3*, and *fea3;bam1d-3* meristems. *bam1d-3* meristems were shorter and narrower than WT, but *fea3;bam1d-3* meristems were the same size as *fea3* meristems. n≥9. p-values calculated using a Student’s t-test. * indicates p<0.05, ** indicates p<0.01, *** indicates p<0.001, n.s. not significant.

**Supplementary Table 4.**
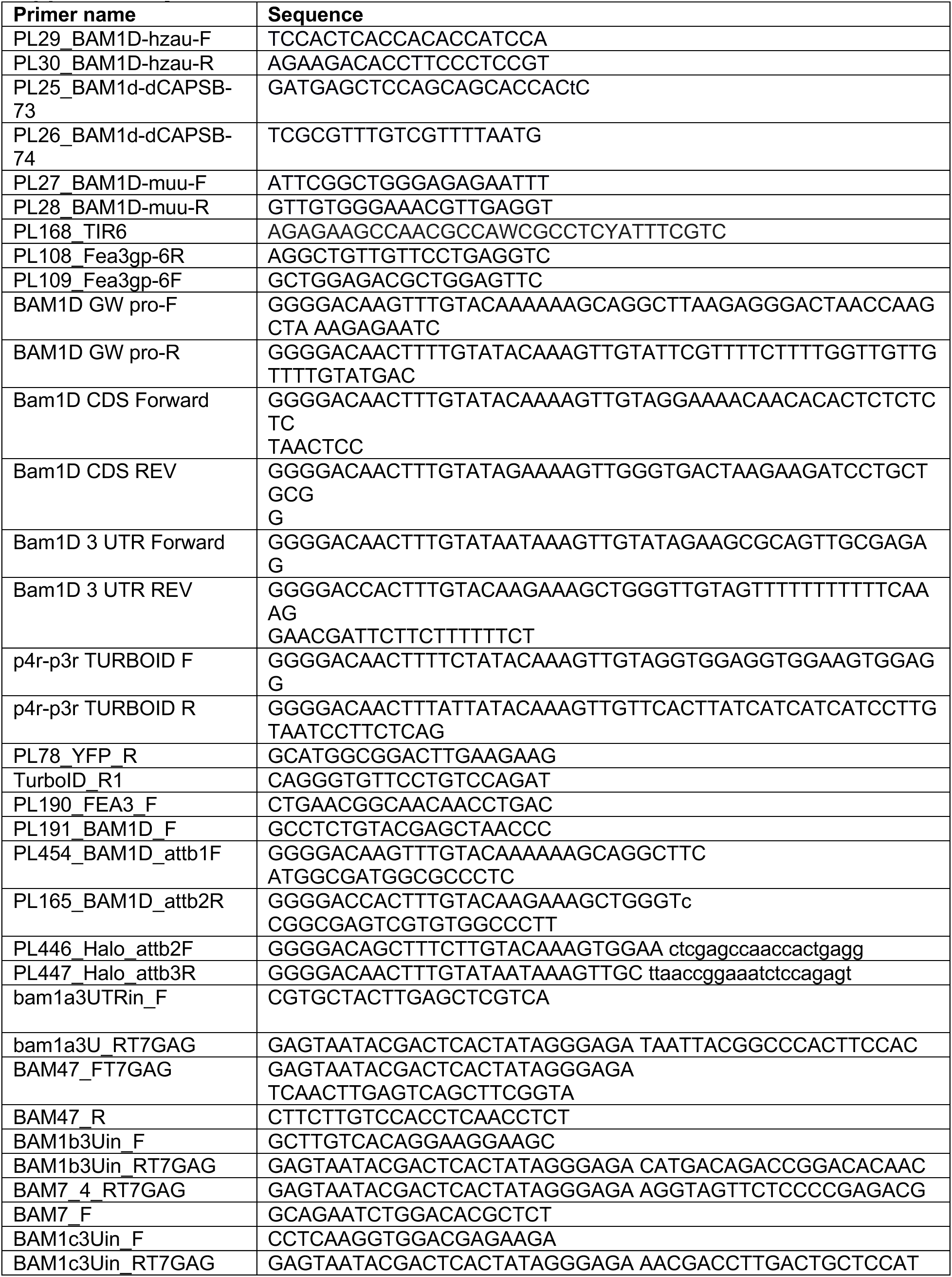
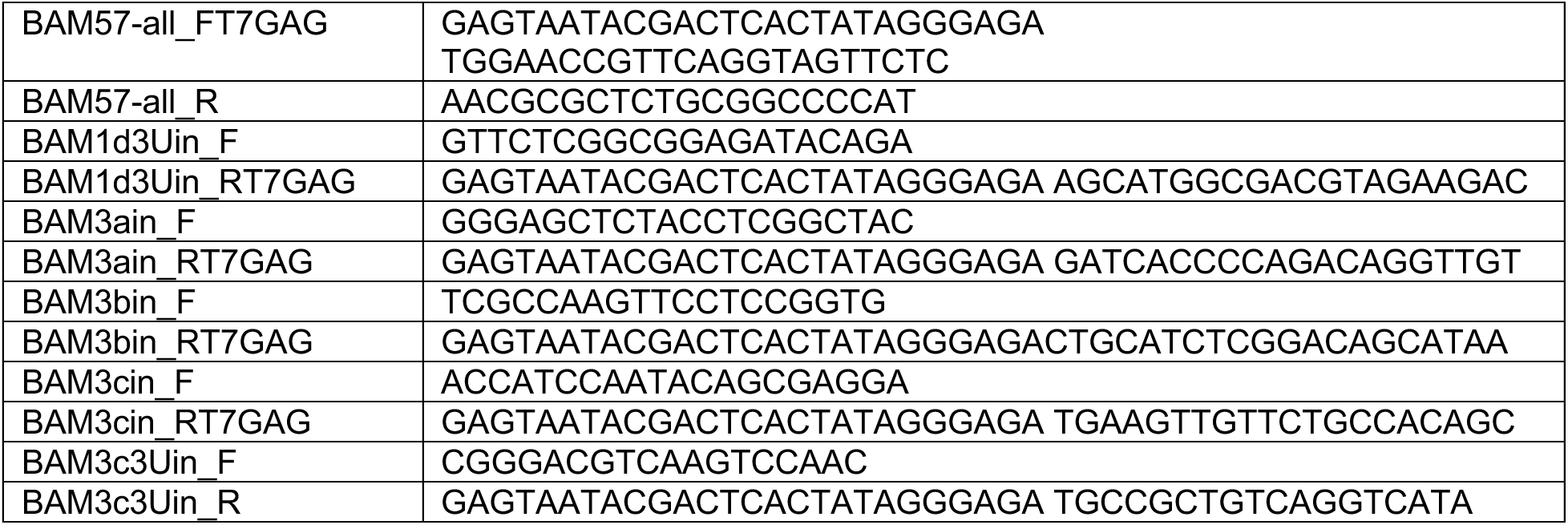
Primers.

**Supplementary Table 5.**
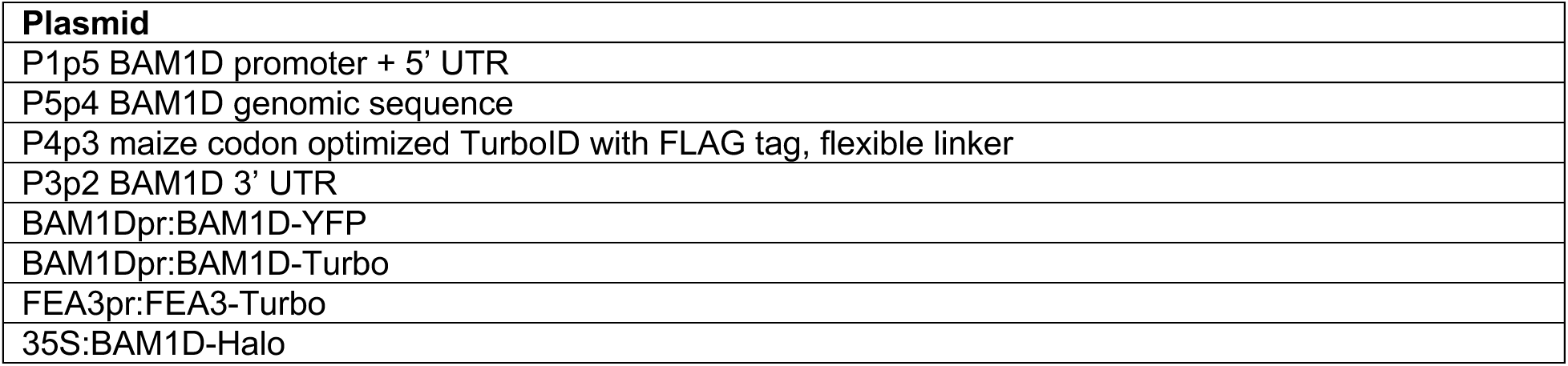
Plasmids.

**Supplementary Table 6.** Peptide sequences.

| Name | peptide | gene ID |
| --- | --- | --- |
| ZmCLE7 | RAVPGGPDPLHH | GRMZM2G372364 |
| ZmFCP1 | REVPTGPDPIHH | GRMZM2G165836 |
| Scrambled peptide/sCLV3 | PPTRGLSHHPVD | NA |
| D15 peptide | PPPQVPSRPNRAPPG | NA |

